# Saturation-seq integrates single-cell saturation genome editing and RNA-seq to quantify *NFE2L2* (NRF2) variant effects

**DOI:** 10.64898/2026.06.30.735631

**Authors:** Magdalena E. Strauss, Andrew J. Waters, Holly Roberston, Timothy Brendler-Spaeth, Aleksander Gontarczyk, Prashant Gupta, Shikha Kataria, Daniel Gitterman, Tumisang Ntereke, Leah Wells, Jamie Billington, Andrew Bassett, Sarah Cooper, David J. Adams

**Affiliations:** European Bioinformatics Institute, Hinxton, Cambridgeshire, CB10 1SD, UK; Department of Mathematics and Statistics, University of Exeter, Exeter, EX4 4QF, UK; Wellcome Sanger Institute, Hinxton, Cambridgeshire, CB10 1SA, UK; Open Targets, Wellcome Genome Campus, Hinxton, Cambridge, CB10 1SA, UK

## Abstract

Interpreting the functional consequences of variants remains one of the central unsolved problems in genomics and clinical genetics. Compounding this, most existing approaches rely on reductive, one-dimensional proxies such as cell growth to score variant effects, which can be a poor substitute for the rich, multidimensional phenotyping that is ultimately needed to understand how variants alter biology. This is especially true for variants known to act through gain-of-function/neomorphic mechanisms. We developed Saturation-seq, a high-throughput platform that combines saturation genome editing with single-cell DNA and RNA profiling to systematically map variant effects. Using CRISPR-based editing in a barcoded haploid cell line, we install hundreds of variants directly into endogenous genomic loci, testing them in multiplex and preserving the native coding and regulatory context. Single-cell amplicon and transcriptome sequencing enables direct linkage of each genomic edit to its transcriptional impact. We apply Saturation-seq to comprehensively characterize 230 variants in the recurrently mutated N-terminal region of *NFE2L2* (NRF2), a master regulator of oxidative stress and an oncogene mutated in lung cancer. We define variant function with ‘disruption scores’ computed from misregulation of known NRF2 targets in single-cell transcriptomes; scores separate pathogenic/benign truthset variants with >90% accuracy and enabled interpretation of TCGA and TRACERx patient tumor data, as well as a rare *NFE2L2* germline variant linked to a developmental syndrome. Thus, we establish a broadly applicable high-resolution single-cell variant-to-function platform with a rich phenotypic readout.

## Introduction

CRISPR/Cas9 loss-of-function (LoF) screens and the CRISPR-based technologies of base and prime editing have transformed our ability to understand and treat genetic disease^1,2^. This is particularly true of conditions associated with recessive disorders, such as tumor suppressor syndromes, where disruption of the disease gene is often associated with cell viability/fitness phenotypes in *in vitro* cell models. Interpreting the functional consequences of genetic variants in oncogenes, which operate via gain-of-function (GoF) mechanisms, remains a further challenge in cancer functional genomics, particularly as clinical sequencing efforts continue to uncover extensive allelic diversity across the driver gene landscape.

Saturation genome editing (SGE) uses Cas9 and Homology Directed Repair (HDR) template libraries to install collections of mutations into the genome, quantifying variant effects as fitness changes in cells through deep amplicon sequencing^3^. This has proven invaluable for LoF variants in essential genes including *BRCA1*, *BAP1*, *VHL*, *RAD51C*, and *DDX3X*, and the non-coding RNA *RNU4-2*, and has been extended to GoF effects in *CARD11*—demonstrating broader applicability^3–9^. However, fitness-based readouts usually require variants to manifest cytostatic or cytotoxic phenotypes, precluding analysis of the broader variant landscape. Critically, fitness alone cannot resolve mechanistically distinct allelic series such as the apoptotic versus cell-cycle-arrest responses documented for *TP53*, nor report on pathway-specific/physiologically relevant molecular phenotypes for GoF/neomorphic, or non-essential LoF variants^10^. These limitations are especially consequential for oncogenes where activating variants may exert pleiotropic, context-dependent transcription-level effects that are often uncoupled from proliferation. Single-cell RNA sequencing (scRNA-seq) provides a clear solution to these limitations by capturing high-dimensional and cell-resolved transcriptional phenotypes^11^. CRISPR-based perturbation screens have successfully leveraged scRNA-seq in pooled formats to unravel gene regulatory networks and drug response mechanisms, including Perturb-seq and CRISPR droplet sequencing (CROP-seq)^12,13^. However, these techniques are often limited to gene-level modulation using CRISPRko, CRISPRi or CRISPRa, with sgRNA sequences used as a proxy for the perturbation outcome^14^. Indeed, to interrogate specific variant-level genomic effects with high-content readouts such as scRNA-seq, other methods of editing – e.g. base-editing, prime-editing or HDR – are required. However, successfully genotyping introduced edits and linking variants to phenotypes is extremely challenging. Functional genomics screens are commonly conducted in multiplex, which further complicates genotype-phenotype linkage, and the challenge is greater still when genome editing is inefficient (e.g. current prime-editing approaches) or introduces unintended mutations (e.g. bystander mutations in base editing) rendering sgRNA identity/detection a poor proxy for detecting cells with a specific/engineered genomic change^15^.

One approach to link genotype to phenotype is through inference of genotype from direct sequencing of transcribed RNA (e.g. GoT-seq, TISCC-seq) but this is insensitive to exonic Single Nucleotide Variants (SNVs) that trigger nonsense mediated decay (NMD) and cannot be used for non-coding variants that do not alter transcribed sequence^16,17^. Methods combining targeted DNA genotyping with scRNA-seq (e.g. Target-seq, CRAFT-seq) rely on low-throughput plate-based workflows, limiting scalability^18,19^. Higher-throughput split-pool and droplet approaches either depend on low-coverage whole-genome DNA sequencing, which is insufficient for accurate single-locus edit calling (e.g. sci-L3_RNA/DNA, DEFND-seq), or only provide targeted RNA profiling, preventing unbiased transcriptome-wide analyses (e.g. SDR-seq, STAG-seq)^20–23^. Building on our previously developed droplet-based framework for linking targeted DNA genotyping of base-edits with whole-transcriptome scRNA-seq via shared cellular barcodes (scSNV-seq), we have developed an approach to enable single-cell transcriptomic readouts for cells with variants precisely engineered into the genome via HDR^24^. This method, called Saturation-seq, is a high-throughput technique and biostatistics platform that combines saturation genome editing with scRNA-seq to comprehensively profile the transcriptomes of specific-coding variants at target regions in multiplex. Saturation-seq enables the assignment of transcriptional phenotypes to specific genomic variants through non-limiting barcodes that are detected in both the DNA and RNA of single cells. This results in molecular phenotype identification with single-cell precision, permitting quantitative (and qualitative) classification of variant function including those that are wild-type-like, LoF (null or hypomorphic), and GoF (hyper or neomorphic), in a way not previously possible.

We apply Saturation-seq to the master regulator of oxidative stress *NFE2L2* (encoding NRF2), a gene recurrently mutated in aggressive, often chemotherapy-resistant, lung squamous cell carcinomas and other tumor types^25^. Activating mutations in *NFE2L2* often cluster in the DLG and ETGE protein motifs, disrupting KEAP1-mediated degradation of NRF2 in the cytoplasm and leading to constitutive, ectopic nuclear-localization and activation of pro-survival/stress-response genes. Accurate variant effect quantification in this context is particularly challenging as not only can GoF effects be pleiotropic, but variants that strongly alter NRF2 function will likely promote cell growth and survival, which will distort population-level (pooled) readouts, thus requiring single cell analyses.

Of note, over 200 transcriptional targets of NRF2 have been identified and are characterized by the presence of antioxidant response element (ARE) 5’ regulatory motifs^25,26^. At these loci NRF2 binds to a sub-sequence (5ʹ-TGACNNNGC-3’) together with dimerized sMAF proteins to induce gene transcription^27,28^. Collectively, these NRF2 target genes are implicated in a wide range of pro-survival cellular processes, including detoxification, redox homeostasis, and metabolism^25^. From these genes we curated a subset of high-confidence targets to report on NRF2 variant effects^29^. Analysis in this way led to the development of ‘disruption scores’ for 230 variants (including 173 missense variants). Importantly, these scores were highly correlated with expected behaviours for clinically significant variants, with 84% sensitivity, 100% specificity and >90% AUC values obtained in Receiver Operating Characteristic (ROC) analysis. Furthermore, disruption score agnostic clustering/dimensionality reduction identified distinct subsets of novel NRF2-activating missense variants, enabling prospective functional classification/assessment of variant effects. We find that disruption scores derived from Saturation-seq data are highly correlated with the transcriptional signatures of ‘the cancer genome atlas’ (TCGA) tumors (vs. matched normal) carrying the same variants, which indicates that HAP1 cells provide a relevant cell-context in which to examine NRF2 function^30^. In addition to variant scoring, we provide differential expression profiles and identify key protein residues with position-based intolerance to variant effects. Crucially, this approach can distinguish different transcriptional profiles even among variants at the same residue.

Our results reveal previously uncharacterized dimensions of NRF2 functional diversity and underscore the need for multidimensional variant effect mapping in the era of precision medicine. More broadly, this work establishes a generalizable technical and analytical framework for the functional interrogation of oncogenic and GoF variation at single-cell resolution, with broad applications in variant interpretation, personalised medicine, precision therapeutic targeting, and multiplexed genetic screening.

## Results

### HAP1 cells exhibit canonical gain-of-function NRF2 phenotypes

HAP1 cells (Horizon Discovery, HZGHC-LIG4-Cas9) have previously been optimized for use in SGE, through introduction of a Cas9 transgene and clonal screening for high Cas9 expression and activity^4^. In addition, these cells were sorted for high haploidy to allow variant effect measurements to be conducted in the absence of potentially un-edited wild-type alleles, which is particularly important for LoF variant assessment.

The mechanism of action for NRF2 regulation is known: KEAP1 binds to NRF2 at DLG and ETGE motifs within the Neh2 domain, which then recruits CUL3 to poly-ubiquitinate NRF2, targeting the protein to degradation within the proteasome (**Fig.1a**)^31–33^. Variants that disrupt KEAP1 binding, particularly those within DLG and ETGE motifs, prevent proteasomal homeostatic regulation on NRF2 in the cytoplasm and allow the protein to enter the nucleus where it transcriptionally activates pro-survival/antioxidant protective genes. Many of these genes are known targets of NRF2 and over-expression can be used as a biomarker for NRF2 activity^29^. To confirm that NRF2 is physiologically regulated and functional in HAP1 cells we performed a series of experiments using an engineered HAP1 cell line carrying a 7bp frameshifting deletion in exon 2 of *KEAP1* (KEAP1Δ, Horizon Discovery HZGHC003774c009), which prevents KEAP1-mediated NRF2 inhibition, phenocopying NRF2 GoF N-terminal mutations. KEAP1Δ cells (clonally expanded) exhibited an increase in NRF2 and an increase in NQO1, a key target of NRF2 transcriptional activation, as measured by western blotting (**Fig.1b**). In addition, KEAP1Δ cells displayed increased fitness compared to wild-type HAP1 cells in colony forming assays (**Fig.1c**). Importantly, platinum-based chemotherapy compounds such as cisplatin induce cell death through DNA damage and reactive oxygen species (ROS) generation^34^. NRF2 pro-survival targets are known to protect against DNA damage and ROS effects^29^. Consistent with this, KEAP1Δ HAP1 cells show approximately two-fold greater viability than wild-type HAP1 cells across physiologically relevant cisplatin concentrations (0-8µM). By MTS assay (**Fig.1d**), the IC50 was 2.326µM (95% CI 2.096-2.582) in KEAP1Δ cells versus 1.408µM (95% CI 1.223-1.620) in wild-type. Alamar Blue gave concordant results (**Fig.1e**): IC50 2.437µM (95% CI 2.237-2.656) versus 1.393µM (95% CI 1.204-1.612). Together these data confirm that HAP1 cells mount a functional NRF2 response.

To confirm that NRF2 variation also elicits detectable fitness and molecular phenotypes in HAP1 cells we next used genome editing to generate an allelic series of clonal lines. Recurrently mutated residues identified in MSK-IMPACT (a clinical tumor sequencing resource) were chosen for installation into the genome; D29, G31, E79 and T80 (**Fig.1f**)^35^. Isogenic/clonal lines containing an inframe deletion of the triplet codon motif D29-L30-G31 (DLGΔ) and the missense variant E79Q (within the ETGE motif) were generated and analysed by MTS assay. Of note, these mutant lines (E79Q and DLGΔ) showed an inherent proliferative advantage independent of drug challenge suggesting enhanced fitness (**Fig.1g**). KEAP1Δ, T80K, DLGΔ and E79Q all showed increased transcription of the NRF2 targets *AKR1C1, NQO1* and *GCLM* by RT-qPCR, with a more modest effect on *GCLC*; notably T80K did not upregulate *GCLC* (**Fig.1h**). Intriguingly, G31R had a limited transcriptional response in terms of mean Log2 Fold-Change (LFC) compared to HAP1 wild-type cells, however *AKR1C1* was moderately increased. As expected, D29*, a LoF variant, did not result in increased transcription, indeed *NQO1* was decreased, suggesting upstream ARE binding by NRF2 is particularly important for the transcription of this gene in HAP1 cells (**Fig.1h**).

We next sought to examine how the transcriptome profiles of clonal variant lines differ. Dimensionality reduction of RNA-sequencing reads showed that KEAP1Δ and DLGΔ transcriptomes were definitively different from HAP1 wild-type, representing the most extreme molecular phenotype we observed. We also analysed (E79Q, G31R, T80K and D29*) (**Fig.1i**). Importantly, missense GoF variants E79Q, G31R, T80K, show distinct transcriptomic profiles from the LoF variant D29*, together with more subtle differences from one another and wild-type, as demonstrated by clear cluster demarcations within a Uniform Manifold Approximation and Projection (UMAP) projection in which KEAP1Δ and DLGΔ were excluded (**Fig.1j**). RNA-seq demonstrates that the variant clonal lines (G31R, T80K, E79Q and DLGΔ) and KEAP1Δ cells exhibit increased expression of known NRF2 targets detected *a priori* of NRF2 association (**Fig. 1k**). As above, NRF2 target genes have well defined roles in detoxification and cellular homeostasis in response to cellular stress^29^. In keeping with this, DLGΔ and KEAP1Δ show increased expression of many of these targets, including those involved in ‘Phase I – drug redox and hydrolysis’ acting to solubilize harmful species/compounds, ‘Phase ∥ – drug conjugation’ of Phase I products to neutralize harmful effects (*UGT1A6*) and ‘Phase ∥ – drug efflux’, to export de-toxified species/compounds from cells (*ABCC3* and *ABCB6*). Cell lines with E79Q and T80K variants showed increased expression of Phase I: *AKR1C1/2/3* & *CES1*, Phase II: *UGT1A6* and Phase III: *ABCB6* enzymes. G31R shows no increase in Phase I or Phase III gene expression, however *UGT1A6* was increased. In addition to these response phases, NRF2 also mediates H_2_O_2_ scavenging through the expression of glutathione-dependent peroxide-reduction enzymes, such as *GPX2*, which has increased expression across all lines, except for the LoF line D29*. These data (**Fig.1**) further demonstrate that HAP1 cells are an appropriate model in which to capture and investigate GoF transcriptomic responses of NRF2 variants.

**Figure 1:**
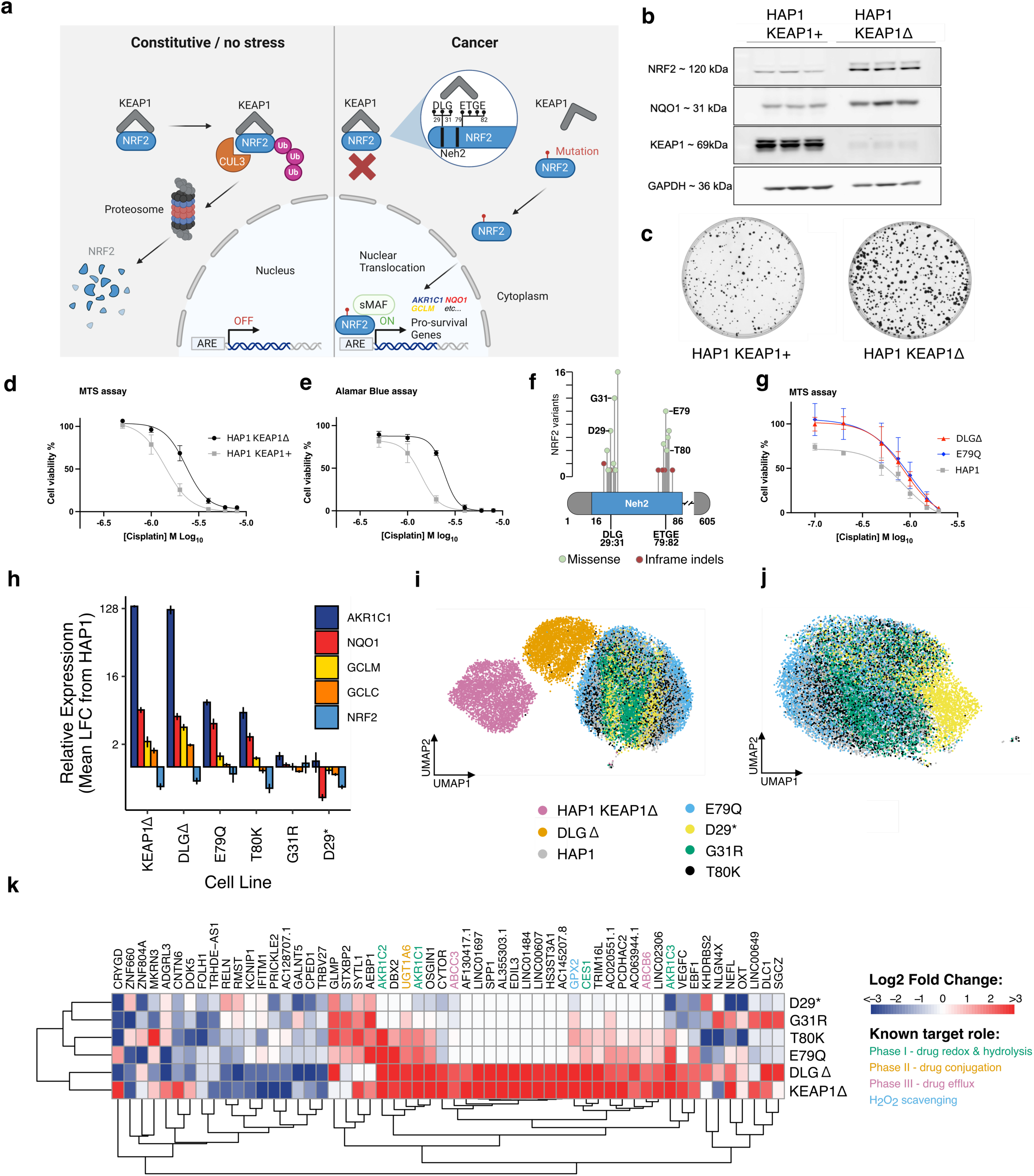
HAP1 cells exhibit canonical gain-of-function NRF2 phenotypes. **a)** Mechanism of action in normal and cancer cells. **b)** Western blot shows that a loss-of-function (LoF) deletion in *KEAP1* (KEAP1Δ) results in increased NRF2 protein levels consistent with the feed-forward effects of KEAP1 on *NFE2L2/*NRF2 expression. NQO1, a key target of NRF2-mediated transcriptional regulation, is also increased. **c)** KEAP1Δ cells have enhanced proliferation compared to wild-type HAP1 cells. In petri dishes 800 cells of each genotype was seeded and grown for 7 days before being stained with crystal violet. This experiment was performed 6 times with similar results, representative examples shown. **d)** KEAP1Δ cells are more resistant to cisplatin than wild-type cells (72hrs exposure in range 0 to 8µM) as measured by MTS metabolic activity assay (2hr exposure to [3-(4,5-dimethylthiazol-2-yl)-5-(3-carboxymethoxyphenyl)-2-(4-sulfophenyl)-2H-tetrazolium]). Media corrected values shown, n=5,000 cells seeded in 96 well plate then cultured for 24hrs pre-exposure, measured in duplicate, error bars +/-SD. Log_10_ cisplatin molar concentration (M) plotted. **e)** KEAP1Δ cells are more resistant to cisplatin than wild-type cells as analysed by Almar Blue assay with 4hr incubation pre-measurement, media corrected values shown, conditions and units as in ‘d’. **f)** Key recurrent somatic mutations reported in MSK-IMPACT are seen at two hotspots within the Neh2 domain of NRF2. Peptide positions of D29*, G31R, E79Q and T80K, analysed in the cell line validation experiments, are highlighted. **g)** Clonal variant lines with a deletion of D29-L30-G31 (DLGΔ) or engineered to carry the E79Q variant are more resistant to cisplatin than wild-type cells as measured by MTS assay in triplicate. Media corrected values shown, conditions and units as in ‘d’. **h)** Real Time-qPCR of clonal line cell cDNA shows that known gene targets of NRF2 are mis-expressed compared to wild-type HAP1 cDNA. Bars show the mean of three biological replicates, lines show +/- standard error of the mean. RT reactions performed using 50ng RNA extracted from confluent wells (6 well plate). **i)** Clonal lines pooled in equal proportions show distinct dimensions (visualised by UMAP) following single cell 10x Chromium transcriptome sequencing and analysis. 1.5e+6 cells per clonal line were cultured and analysed on a 10x Chromium machine. DLGΔ and KEAP1Δ produced more profound transcriptional phenotypes than single nucleotide variants; E79Q, D29*, G31R and T80K. **j)** The LoF variant D29* shows a distinct transcriptional signature (visualised by UMAP) when compared to likely GoF variant lines; E79Q, G31R and T80K. E79Q appears more distinct from wild-type HAP1 than G31R and T80K. **k)** Heatmap of gene expression in clonal lines compared to wild-type shows differing profiles between lines. DLGΔ and KEAP1Δ show an increase in many known NRF2 targets with defined functions of redox response; Phase I (green), Phase II (orange), Phase II (pink) and hydrogen peroxide scavenging (blue). E79Q and T80K also show over-expression of several of these targets, including Phase I targets *AKR1C1,2,3* and *CES1*. G31R shows over-expression of Phase II and H_2_O_2_ scavenging genes *UGT1A6* and *GPX2*, respectively. Dendrograms show similarity clustering between lines (down) and genes (across), respectively.

### Saturation genome editing in a HAP1 barcoded cell line

To allow the association of DNA genotype and RNA phenotype in single cells (see Methods) we transduced a population of HAP1 HZGHC-LIG4-Cas9 cells with a barcode construct amenable to detection in DNA and RNA^24^. RNA detection is possible due to the incorporation of a barcode downstream of the PGK-promoter driving BFP (**Fig.2a**). To maintain high barcode complexity, 4^10^ unique barcode constructs (1,048,576) were introduced into 18e+6 cells (MOI=0.3, ∼4-5e+6 single integrations) with 8e+6 haploid BFP+ cells sorted for downstream analysis. FACS gates for haploidy and BFP+ sorting are shown in **Extended Data Fig.1a** and **b**, respectively.

Genomic regions within the Neh2 domain of the canonical ‘Matched Annotation from NCBI and EMBL-EBI’ (MANE) *NFE2L2* transcript (ENST00000397062.8) which encodes NRF2 were identified for Saturation Genome Editing (SGE) library design (**Fig.2b**)^36^. These ranges are termed the DLG (GRCh38, Chr2:177,234,247-177,234,218) and ETGE (GRCh38, Chr2:177,234,088-177,234,059) target regions and contain 10 codons each (**Fig.2c,d**). SGE HDR template libraries were generated using VaLiAnT software to include all single SNVs, codon deletions (sequential inframe deletions of each codon), 1bp deletions, a stop-gained scan, and all possible missense changes (multi-nucleotide changes, MNVs) at each codon, resulting in n=333 and n=329 unique oligonucleotide variant containing tracts for DLG and ETGE target regions, respectively^37^. To prevent Cas9 cleavage of successful genomic variant installations, three library-specific synonymous SNVs were incorporated in all oligonucleotide tracts at PAM/Protospacer sites, preventing sgRNA binding (**Fig.2c,d**). Barcoded HAP1 cells were transfected with sgRNAs and HDR template libraries (6e+6 cells per target region) and cultured for 14 days in total (see Methods).

To determine cloning and editing efficiency, we performed next-generation sequencing (NGS) of HDR template library plasmids and edited cell pools (**Extended Data Fig.2**). 100% of desired variants were successfully and uniformly cloned (Gini coefficient = 0.08 and 0.14 for DLG and ETGE regions, respectively) and 100% were successfully installed into the genome (mean coverage of the edited pools was 887x and 432x per variant for DLG and ETGE target regions, respectively, measured in technical duplicate) (**Extended Data Fig.2a,c**). Genome editing occurred with high HDR efficiency (36.2% and 32.6% for DLG and ETGE target regions, respectively). The Saturation-seq workflow (**Fig.3a-f**) requires the same unique barcode to be detected in the RNA and DNA from single cells within the same pool/population of edited cells. We therefore reduced barcode complexity after editing to ∼3,000 per experiment (**Fig. 3b**). Prior work suggests ∼3 cells per barcode gives good genotype recovery in single-cell amplicon sequencing, but anticipating increased cell stress from editing, selection and passaging, we targeted 10 cells per barcode across ∼300 variants per region (see Methods)^24^. In addition to a reduction in complexity, this process also permits the proliferation of multiple daughter cells of the same variant-barcode association, allowing replicate detection (we achieved 1-2 barcodes per variant, with genotyping associations that had ≥3 cells used, see Methods for calculations). Post-bottlenecking, 80% of DLG (n=265/333) and 82% of ETGE region (n=270/329) variants were highly represented (measured as >5% relative counts) within edited cell populations. As expected, an increase in Gini coefficient (from 0.17 to 0.55 for DLG, and from 0.19 to 0.58 for ETGE) and negative binomial distribution (right skew) due to bottlenecking pressure was observed indicating a desired reduction in barcode to variant complexity (**Extended Data Fig.2b,d,e,f**); variant counts were subsequently processed through an informatics pipeline (documented in Methods). No strong positional effect or molecular consequence bias was observed in the installation of SGE variants (**Extended Data Fig.2g,h**). Cells were grown for 14 days post-transfection, to allow for selection, bottlenecking and expansion. To confirm that GoF molecular phenotypes were active over the length of a tissue culture screen we transfected wild-type HAP1 cells with sgRNAs and HDR templates to generate polyclonal DLGΔ and E79Q variant lines, observing that *NQO1* and *AKR1C1* expression was increased from Day 4 until D18 (inclusive of D4, 10, 14, 16, 18) as measured by RT-qPCR for both genome edited cell populations (**Extended Data Fig.2i**).

**Figure 2:**
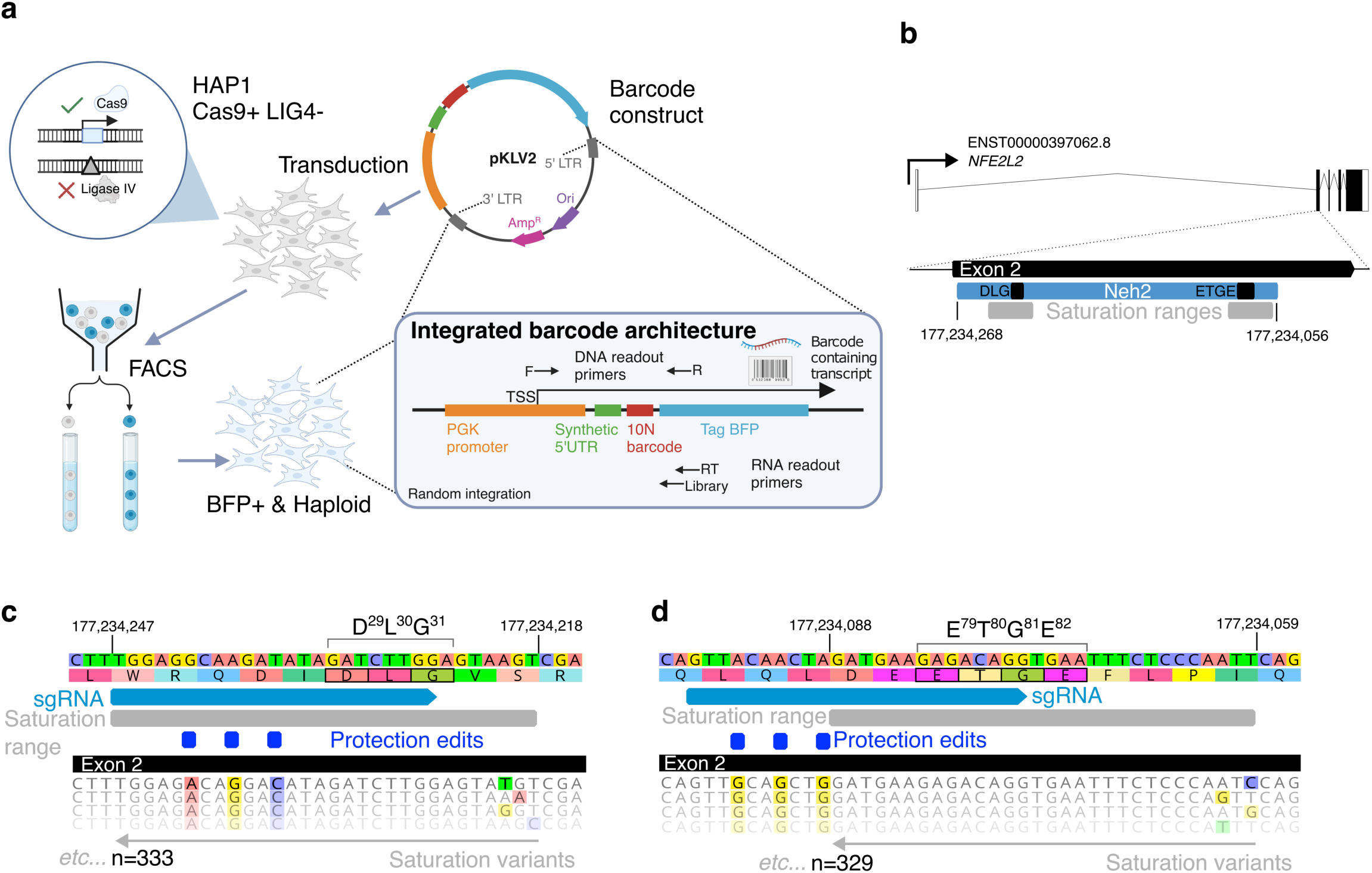
HAP1 cell barcoding and gene editing. **a)** HAP1-A5 cells expressing Cas9 and with a 10bp deletion in *Ligase IV* (*LIG4*, to favour HDR over NHEJ) were transduced with a barcode construct containing a synthetic 5’UTR upstream of a random 10 nucleotide sequence and a BFP reading frame. When integrated randomly in cells the barcode is expressed from the PGK promoter. 8e+6 cells were sorted for positive integrations by BFP+ and for 1N ploidy. **b)** Saturation Genome Editing was performed on regions around the peptides DLG and ETGE within the Neh2 domain of exon2 of the transcript ENST00000397062.8. GRCh38 coordinates are shown. Saturation ranges where variants were installed are shown in grey. **c)** DLG target region in detail. sgRNA binding position is shown in light blue, synonymous protection edits are shown in dark blue, which are fixed changes present in all generated oligonucleotides to prevent Cas9 cleavage of previously installed edits. DLG peptides are highlighted in black boxes around the key amino acids within the NRF2 translation. 333 variants were designed to be included in the saturation range (see Methods). **d)** ETGE target region in detail. ETGE peptides are highlighted in black boxes around the key amino acids within the NRF2 translation. 329 variants were designed to be included in the saturation range (see Methods).

**Figure 3:**
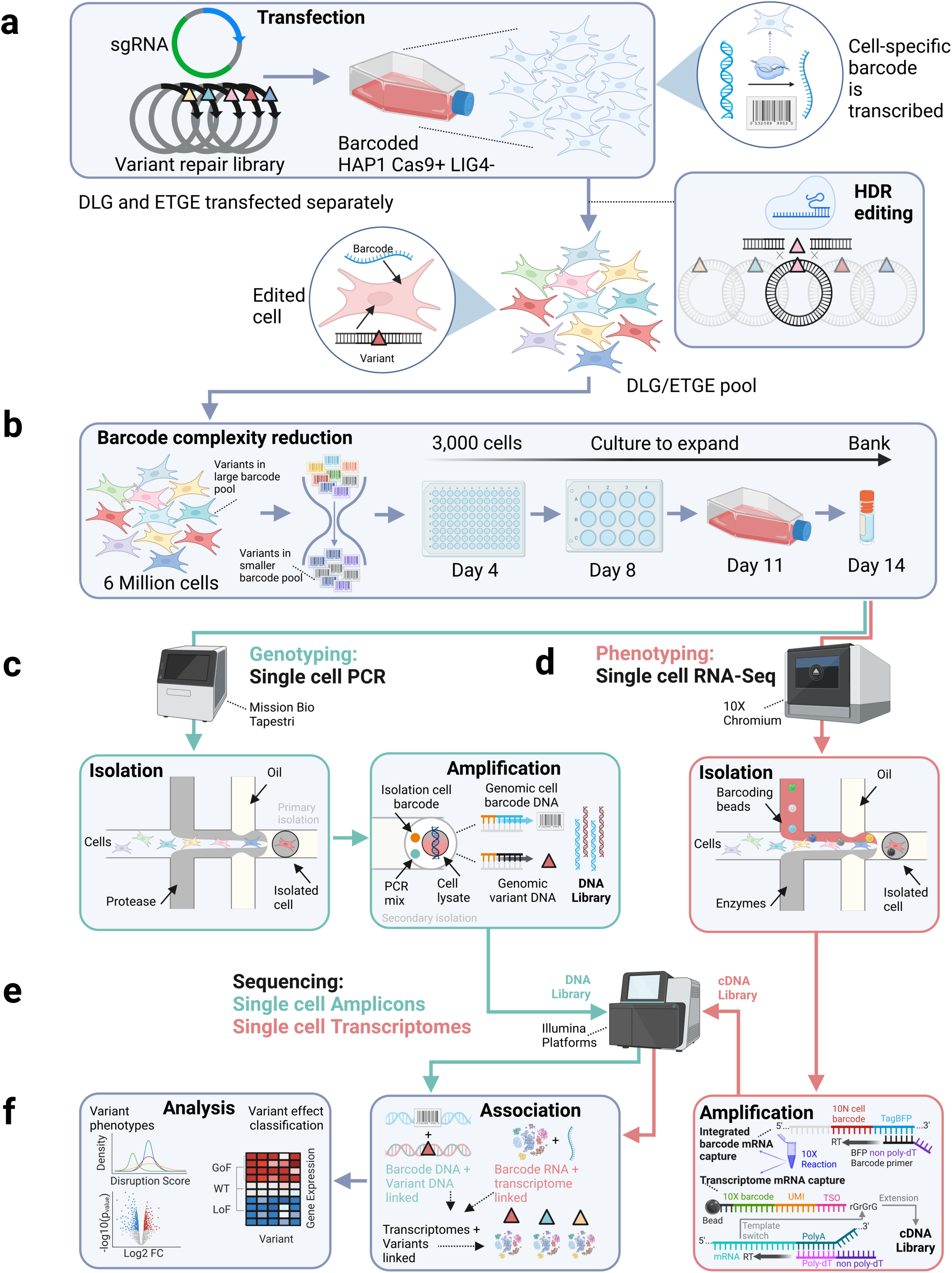
Experimental overview of Saturation-seq. **a)** Loci are edited through transient co-transfection of an sgRNA and cognate HDR repair template library containing variants into barcoded HAP1 Cas9 expressing cells that are null for *LIG4*. Barcodes are 10N DNA sequences that are transcribed to RNA (top right, circle inset), HDR installs variants into the DLG and ETGE hotspots through separate transfections, resulting in cells that contain a variant and a DNA/RNA barcode (top left, circle inset). **b)** To reduce the number of barcodes associated with the same variant, and to permit technical replicates (the same barcode-variant association seen in multiple cells at subsequent stages), of 6 million cells transfected for each of DLG and ETGE pools, 3,000 cells (intended to be ∼10 barcodes per variant, with 1-2 barcodes successfully detected) were passaged at Day 4, and expanded over 14 days of tissue culture. **c)** Single cell PCR: ETGE and DLG edited cells were genotyped using Mission Bio proprietary primer libraries to produce single cell amplicons from edited loci and barcode integrant genomic templates, this process involves the isolation of single cells in microfluidic oil droplets and the provision of PCR reagents and cell-isolation barcode in a secondary, amplification encapsulation (green boxes/arrows). **d)** Single Cell RNA-Seq: the same pools of ETGE and DLG edited cells processed in ‘c’ are isolated/encapsulated in microfluidic oil droplets together with barcoding beads for 10X RNA-sequencing. A 10X amplification of integrated barcode and mRNA in isolated cells is shown (bottom right box, red); a non-poly-dT reverse transcriptase (RT) primer (black/purple) complementary to TagBFP sequence (blue) 3’ of 10N cell barcode (red) permits RT conversion of barcode from RNA to cDNA, and the 5’ end of the barcode transcript template switches onto the 10x barcoded beads (containing the cell barcode and unique molecular identifier (UMI, unique for transcript) in an identical manner to the single cell transcriptome (5’ GEM-X 10x Genomics kit). A UTR barcode specific library is produced from the amplified cDNA using a nested PCR after SPRI cleanup (see Methods). **e)** DNA libraries from single-cell variant and barcode genotyping (green) and cDNA from single-cell transcriptome and barcode libraries (red) are sequenced on Illumina platforms**. f)** Informatic analysis associates DNA variants to their cognate transcriptomes through barcode sequences captured in single cells (bottom centre box), post-associative analysis uses biostatistics to calculate variant-specific disruption scores computed from the degree of transcriptome aberration manifested from the variant, and differential gene expression to identify known and novel variant-specific mis-expression profiles (bottom left box).

### Coupling of DNA variant calls to RNA transcriptomes in single cells

Saturation-seq combines SGE (**Fig.3a,b**) with single-cell sequencing of both DNA and RNA, recovering edits and barcodes from each modality respectively (**Fig.3c-e**), followed by computational linkage of barcodes across the two readouts (**Fig.3f**). This allows single-cell transcriptomes to be assigned to specific, genomically installed variants produced in multiplex, so that each variant is associated with a distinct transcriptomic phenotype. The DLG and ETGE motif hotspot regions were targeted separately using HDR template libraries of 333 and 329 unique variants respectively (**Fig.3a**), comprising in total: 47 synonymous, 28 stop-gained, 506 missense, 46 frameshift (1 bp deletions), 29 in-frame single-codon deletions and 6 multi-codon or serial deletions.

Unless stated otherwise, ‘barcode’ throughout refers to the 10N untranslated region (UTR) barcode (**Fig.2a**, **Fig.3d**), with ‘cell barcode’ referring to droplet distinction barcodes from barcoding beads (**Fig.3d**).

Edited DLG and ETGE cell pools were single-cell genotyped via Tapestri (Mission Bio, ≤15,000 cells/reaction) and transcriptomically profiled with the 10x Genomics 5’ HT method (∼60,000 cells/lane), with a spiked BFP-specific RT primer enabling enriched cell barcode capture; libraries were sequenced on MiSeq (250PE) and NovaSeq 6000, respectively (see Methods). Reads were mapped to amplicons (BWA-MEM, Q≥30), filtered to retain only wild-type or valid SGE editing outcomes, and droplets cleaned for doublets/noise using 90% read-dominance thresholds and Hamming-distance barcode correction, yielding 1,927/1,928 droplets and 217/242 barcodes across 149/154 genotypes for the DLG and ETGE datasets, respectively (see Methods)^38^. Variant recovery was 56.2% (149/265) and 57.0% (154/270) of post-bottleneck complexity for the DLG and ETGE regions respectively, corresponding to 44.8% (149/333) and 46.8% (154/329) of the original library complexity. We then focused on variants detected in both the DNA (Tapestri) and RNA (10x) single-cell datasets, enabling us to link each variant to a transcriptome profile and compute per-variant ‘disruption scores’ (see below, and Methods). In the final score dataframe, we present variants where we were able to compute different forms of disruption score (‘integrated’ and ‘specific’ see below); we also include multi-residue hotspot deletions where only specific scores could be calculated, resulting in a total of n=230 variants; See **Supplementary Table 1**.

### Saturation-seq identifies a spectrum of missense effect correlated with redox ontology

As the data are extremely rich, to observe and appraise trends across genotypes it is necessary to reduce RNA phenotype dimensions. Therefore, we computed a variant effect functional score based on the disruption of a curated list of 82 known NRF2 target genes (derived from an expert review, see Methods for gene list used) across both DLG and ETGE target regions^29^.

Next, we produced both ‘integrated disruption scores’ and ‘specific disruption scores’ for variants, to assess trends across both DLG and ETGE regions, and more sensitive variant effects within regions, respectively. Both are based solely on abundance changes of known NRF2 targets. For both metrics, a positive score indicates increased transcription of key NRF2 targets and a negative score, a decrease. Integrated disruption scores were computed as follows: DLG and ETGE region cell/variant scRNA-seq datasets were integrated based on high confidence known NRF2 target expression profiles using the mutual nearest neighbours method, then to avoid pseudo-replication bias, we calculated the median of the principal component (PC) scores across barcodes for the same variant genotype, finally we calculated the median across genotypes for the same protein-level change (i.e. average of redundant codon changes). Specific disruption scores were computed through the same process without harmonization between region experiments (see Methods). To aid interpretation, both scores were scaled such that the median score for wild-type was 0, and the maximum median score for a group of barcodes sharing one genotype was 1.

Genomic editing events that contain PAM/Protospacer Protection Edits (PPEs, fixed synonymous variants which prevent Cas9-cleavage of installed tracts) but are otherwise wild-type (no additional variants) are expected to have a largely wild-type transcriptional response (**Fig.2c,d**); using PPE only events as a baseline reference rather than GRCh38 wild-type (unedited cells) sequence is expected to normalize for any transcriptional response due to editing in general, ‘wild-type’ refers to ‘wild-type (PPE)’ unless otherwise stated. In addition, when computing wild-type disruption scores we treated individual wild-type barcodes as replicative controls (rather than averaging across all wild-type barcodes), increasing baseline (no effect) sampling.

Integrated disruption scores of wild-type cells (containing only PPE variants) exhibit a unimodal/normal distribution at zero (median=-0.009, n=30 barcodes, n=448 cells), as expected due to median scaling of wild-type scores. Missense variants (median=0.0913, n=2494 cells, n=173 variants [median n=10 cells per variant]) are bimodally distributed/positively skewed, identifying a subset of variants which positively disrupt the transcription of known NRF2 targets (**Fig.4a**). As expected, synonymous variants are centred around 0 and do not have significantly different disruption scores from wild-type (pairwise two-sided Mann-Whitney U test, Benjamini-Hochberg (BH) FDR correction, q=0.104, n=195 cells, n=13 variants [median n=6 cells per variant]); missense variants (n=2492 cells, n=173 variants, as above) and inframe codon deletions (n=336 cells, n=11 variants [median n=6 cells per variant]) have more positive disruption scores compared to wild-type (q=0.026 and q=0.006, respectively), and stop-gained (n=137 cells, n=11 variants) and frameshift variants (n=336 cells, n=19 variants) have more negative disruption scores compared to wild-type (q=0.0002 for both) (**Extended Data Fig.3a, Fig.4b,c**).

**Figure 4:**
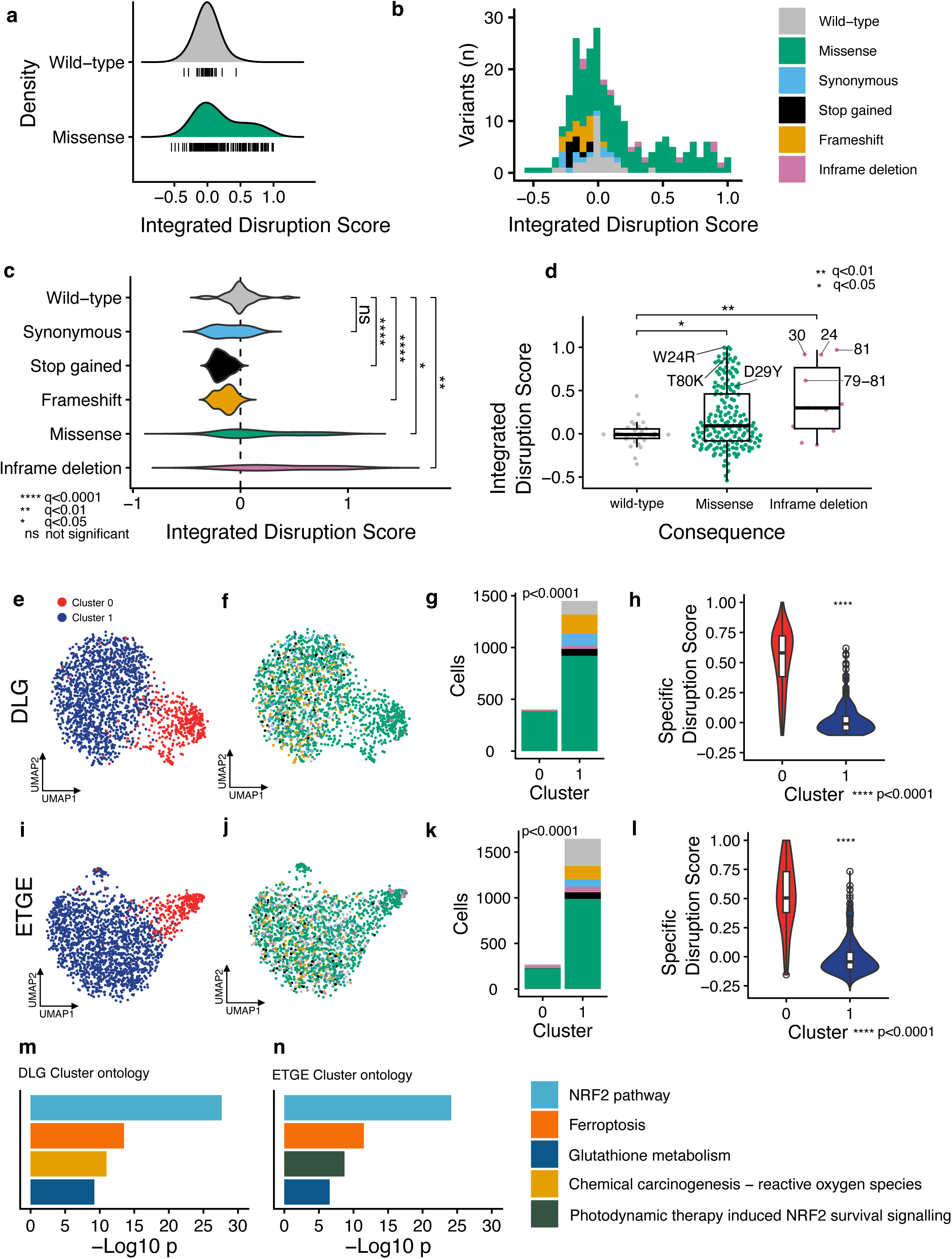
Missense variants demonstrate a spectrum of NRF2 gene network disruption. **a)** An integrated disruption score calculated from changes in known NRF2 targets for variants across the DLG and ETGE target regions is unimodal around 0 for wild-type (median score =-0.009, n=448 cells, n=30 barcodes), whereas missense variants are positively skewed with a bimodal pattern (median score=0.0913, n=2494 cells, n=173 variants [median n=10 cells per variant]), indicating a class of missense variants that are more disruptive to NRF2 target expression. Black tick marks show individual wild-type barcodes and missense variants. **b)** Histogram showing distribution of integrated disruption score coloured by molecular consequence. Missense variants demonstrate a positively skewed/bimodal distribution (cell/variant numbers as in ‘a’), inframe deletions of codons or peptides are observed to have a disruption score centred around 0, or high disruption. As expected for a GoF mechanism, LoF stop gained (n=137 cells, n=11 variants) and frameshift variants (n=336, n=19 variants) do not result in known NRF2 target positive disruption, but do appear to be negatively disruptive, centred below 0. Synonymous variants are centred around 0, indicating that these variants are non-disruptive (n=195 cells, n=13 variants, [median n=6 cells per variant]). **c)** Missense variants and inframe deletions show similar distribution of variant effect, as seen in previous SGE studies and are significantly positively disruptive compared to wild-type (Mann-Whitney U test, BH FDR correction, q=0.026 and q=0.006, respectively), synonymous variants are not disruptive compared to wild-type cells (q=0.104), stop-gained and frameshift variants show significantly negative disruption scores (q=0.0002 for both). **d)** Missense variants have a significantly higher median disruption than wild-type cells (q=0.026). Key recurrent, oncogenic variants (labelled) are observed to be disruptive when installed via multiplexed gene editing, as expected. Inframe deletions are significantly different from wild-type in aggregate (q=0.006) and form two distinct groups, those that are more disruptive are all within key protein positions of DLG and ETGE or recurrently mutated W24. Box plots show interquartile range, horizontal line shows median integrated disruption scores, whiskers show maximum and minimum value, which is not an outlier. **e)** Dimensionality reduction and k-means clustering on n=1,853 cells identify two distinct clusters (0 and 1, coloured red and blue, respectively) of transcriptomic activity for DLG target region variants. **f-g)** DLG target region variants show differing mutational consequences between clusters 0 and 1, with proportionally more missense in Cluster 0, colours as in ‘Fig.4b’ (χ^2^ = 167.41, p<0.0001); DLG target region variants in Cluster 0 are almost exclusively missense variants (g). **h)** DLG target region variant containing cells in Cluster 0 (n=399/1,716 cells plotted [n=137/1,853 cells were not possible to score so are not plotted]) have significantly higher aggregate specific scores than cells in Cluster 1 (n=1,317/1,716 cells), violin plots show distribution of disruption scores calculated for DLG variants, box plots show interquartile range, horizontal line shows median DLG specific disruption score, whiskers show maximum and minimum value, which is not an outlier, Mann-Whitney U test, ****p<0.0001. **i)** as for ‘e’, with ETGE target region, n=1,917 cells. **j-k)** as for ‘f-g’, with ETGE target region (χ^2^ = 115.1, p<0.0001). **l)** as for ‘h’, with ETGE target region. Cluster 0 n=260 out of a total of n=1,606 plotted (n=311/1,917 cells were not possible to score so are not plotted), Cluster 1 n=1,346/1,606. Mann-Whitney U test, ****p<0.0001. **m)** Cluster 0 DLG ontology analysis shows mis-regulation of key NRF2-related biological processes, top 4 ranked by -Log10p are shown. **n)** Cluster 0 ETGE ontology analysis shows mis-regulation of key NRF2-related biological processes, top 4 ranked by -Log10p are shown.

This suggests that GoF and putative LoF transcriptional responses are detectable in aggregate for missense and codon deletions, and stop-gained and frameshift variants of *NFE2L2*, respectively. Inframe deletions of codons show distinct bimodal variant subsets (**Extended Data Fig.3a, Fig.4b, c**) which are present over a dynamic range comparable to that seen for missense variants (as has been seen with previous SGE studies) with deletion of canonical pathogenic variant codons identified as disruptive (**Fig.4c,d and Extended Data Fig.3b**). Canonical pathogenic variants T80K (c.239C>A), W24R (c.70T>A) and D29Y (c.85G>T) are all classed as disruptive by Saturation-seq, with integrated disruption scores of 0.8714, 0.9966 and 0.5480, respectively (**Fig.4d**). T80K was also assessed as disruptive at the clonal line level (**Fig.1**). Moreover, genes identified to be mis-regulated by differential gene expression in clonal lines (represented in **Fig.1k**) are also found to be mis-regulated in single-cell transcriptomes across a variety of variants in both the DLG and ETGE regions, including increased expression of key NRF2 targets such as *AKR1C1/2/3* and *GPX2* (**Extended Data Fig.3c,d**).

Unlike targeted approaches, Saturation-seq captures whole transcriptomes, permitting unbiased discovery of variant-associated transcriptional behaviours beyond known NRF2 targets. K-means clustering on Saturation-seq processed cell populations (see Methods) identified discrete cell subsets with transcriptional profiles that correlate with molecular consequence and known target mis-expression (**Fig.4e-l**). For both the DLG and ETGE target regions, a distinct NRF2-disrupted ‘cluster 0’ population was apparent (n=403/1,853 cells for DLG, n=271/1,917 cells for ETGE; **Fig.4e,i**). Importantly, Cluster 0 cells are composed of predominantly missense and inframe deletions for both DLG (98.0%, n=395/403, **Fig.4f,g**) and ETGE (92.6%, n=251/271, **Fig.4j,k**), additionally, compared to Cluster 0, Cluster 1 contains proportionally more cells edited to contain stop-gained and frameshift variants (for DLG: 4.5%, n=65/1,450 in Cluster 1, versus 0%, n=0/403 in Cluster 0 and for ETGE: 4.25%, n=70/1,646 in Cluster 1, versus 0.74%, n=2/271 in Cluster 0 for ETGE); this is consistent with cluster 0 cells being differentiated from cluster 1 cells through GoF NRF2 transcriptional activity. Indeed, DLG and ETGE Cluster 0 cells have significantly increased expression/positive disruption of known NRF2 target genes compared to Cluster 1 cells in each region, as evidenced by significantly higher ‘specific disruption scores’, confirming GoF NRF2 transcriptional activity (two-sided Mann-Whitney U test, p<2.2e-16, for DLG and ETGE), (**Fig.4h,l**). Furthermore, Metascape ontology analysis, that is blind to disruption scoring/curated lists of known target mis-regulation, identified Cluster 0 cells across DLG and ETGE regions to be significantly enriched for ‘NRF2 pathway’ components along with other NRF2-related functional ontologies, such ‘Glutathione metabolism’ and responses to reactive oxygen species (**Fig.4m,n**).

### Missense variants exhibit positional and transcriptomic effects over key oncogenic residues

The DLG (amino acids 29:31) and ETGE (amino acids 79:82) motifs are the canonical oncogenic hotspot residues within the Neh2 domain of NRF2 and contain 42.1% n=273/648 of recurrent mutations (>1 accession in MSK-IMPACT, n=648/944 *NFE2L2* variants, April 2026) (**Fig.1f**). To interrogate these motifs and to investigate the oncogenicity of flanking residues, we assayed amino acid residues 24:33 and 77:86 of the DLG and ETGE domains, respectively. Specific disruption scores show a correlation with peptide position for DLG and ETGE regions (**Fig. 5a-d**). Missense variants in the DLG region exhibit a high density/concentration of positive disruption scores across the canonical 29:31 range, together with W24, an additional recurrently mutated residue, with non-recurrent R25 and S33 showing a greater density of score values around zero (**Fig.5a,b**). ETGE has a very clear positional effect, with minimal disruption seen at non-canonical residues 83:86 and positive disruption at 79:82, together with D77 an additional recurrently mutated residue (**Fig.5c,d**). The more discrete positional effect seen in ETGE relative to DLG is consistent with the known higher binding affinity of the ETGE motif for KEAP1 compared to DLG^39^. Positional effects can also be seen at the single-cell transcriptomic level–as measured by *de novo* detection of differentially expressed genes, (i.e. not based on clonal line detected mis-regulation as is the case in **Extended Data Fig.3c,d**)–where R25 and S33, and F83:I86 variants are mostly resistant to GoF effects, with specific variants at other amino acid positions exhibiting degrees of increased expression for known NRF2 targets in addition to other genes; additionally, different transcriptional profiles are seen between different putatively activating variants (**Fig.5e,f**).

**Figure 5:**
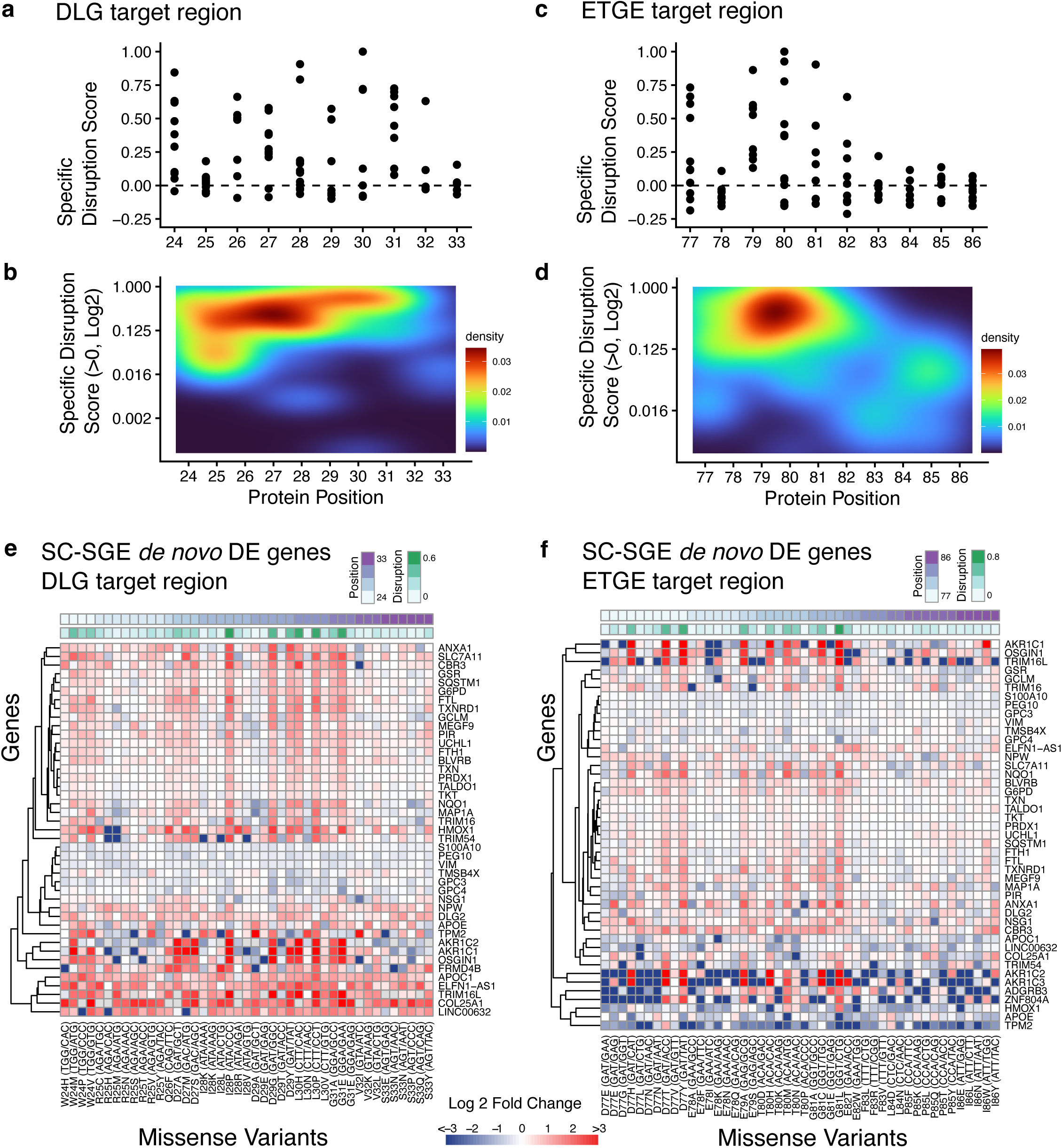
Missense variants are highly disruptive at canonical motifs. **a)** Missense variants across the DLG target region show a dispersed pattern. Variants with high specific disruption scores (likely GoF transcriptional effect) cluster around the DLG protein positions (29:31). **b)** The same data in ‘a’, shown as a density plot, specific disruption scores for individual missense changes are collectively higher at the DLG motif and W24 (a known, recurrently mutated codon) than at other adjacent protein positions assayed, notably positions 25 which has an increased density at lower scores, and 32 and 33 have a spread of lower scores. **c)** The ETGE region shows higher disruption scores at positions 79:82, the key ETGE positions. Interestingly position 77 is also intolerant to missense change. 83:86 have collectively lower disruption indicating missense changes are extensively tolerated at these positions. **d)** Density plot of disruption score for ETGE shows missense variants at position 77, 79:82 are similarly highly disruptive. A cluster of density is seen at the bottom right for variants above 0 that are similarly low in disruption score for positions 83:86. **e)** Heatmap coloured by LFC of gene expression from wild-type, showing genes found to be differently expressed in missense variants within the DLG target region compared to wild-type (discovered *de novo*). Clustering of gene expression shows key NRF2 targets are increased in expression at canonical motif positions as well as additional protein positions. Missense variants ordered by increasing protein position. **f)** As e but for the ETGE target region. A very clear positional effect of GoF variants is seen.

### Missense variant transcriptomes in HAP1 are highly correlated with tumor RNA profiles

HAP1 cells were characterized to have detectable NRF2 GoF molecular phenotypes (**Fig.1**). However, to further investigate cell-context in terms of physiological/disease-relevance of the model and the Saturation-seq methodology we compared the single-cell transcriptomic profiles of variants assayed by Saturation-seq that were also found to have matched tumor-normal samples in TCGA (**Fig.6a-d, Extended Data Fig.4**).

**Figure 6:**
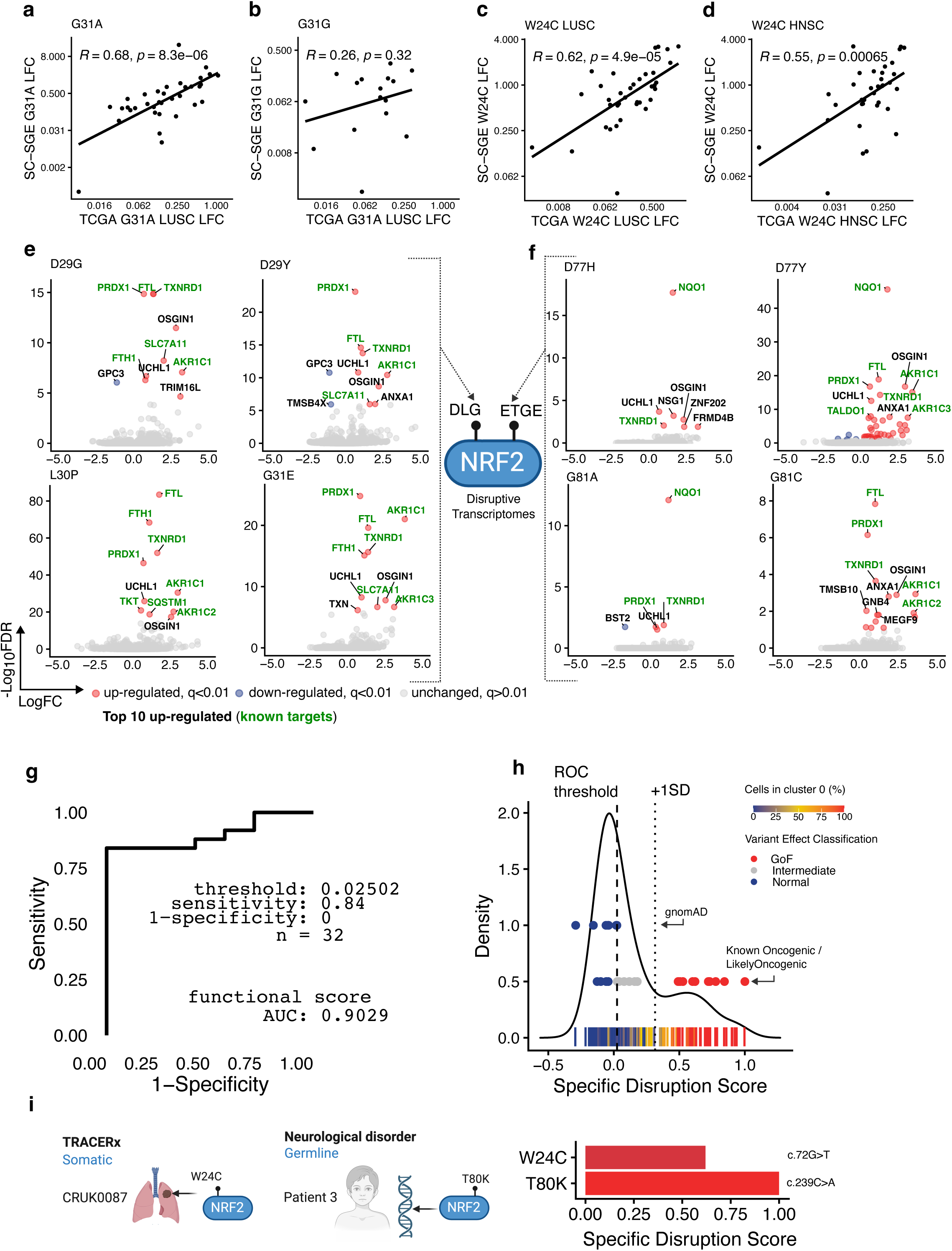
Clustering reveals distinct missense variant classifications that aligns with clinical somatic variant data. **a)** SC-SGE LFC compared to TCGA matched-normal tissue sample LFC for G31A LUSC missense variant containing tumor shows highly correlated effects for positive LFCs for known NRF2 targets. **b)** A significant positive correlation is not seen for SC-SGE LFC of genes in G31G, a synonymous variant at the same residue. **c-d**) Similar to ‘a’ W24C tumors also show positive correlations between Saturation-seq derived SC-LFCs and TCGA LFCs, for LUSC (c) and HNSC (d). **e)** Differential gene expression analysis of Saturation-seq using GLMMs (Methods) derived single-cell transcriptomes for recurrent variants at each of D29, L30 and G31 of the DLG motif show significant, known (green, curated list derived from Robertson *et al*. 2020), and other/unknown (black) target mis-regulation (top 10 by decreasing -Log10FDR displayed) **f)** Similar to ‘e’ mis-regulation of known (curated list) and other/unknown targets are seen at exemplar positions flanking/within the ETGE motif. **g)** ROC curve calculated with 25 true positives (MSK-Impact recurrent GoF missense variants with oncogenic or likely-oncogenic status, n=23 missense, n=2 inframe deletions) and 7 true negatives (expected to be non-compromising or likely-/benign, n=4 synonymous, n=2 missense, n=1 stop-gained). A high classification accuracy of >90% is seen when using DLG and ETGE specific disruption scores. Specific disruption scores for NRF2 are seen to be highly sensitive (84%) and specific (100%) at the ideal score threshold (0.02502). **h)** A bimodal distribution of DLG/ETGE specific disruption scores is seen for n=230 genotypes/variants. The specific disruption score threshold at which maximum sensitivity/specificity is seen in ‘g’ is shown as a vertical dashed line (x=0.02502), dysfunction and non-dysfunctional variants are separated at this threshold, with 1 positive standard deviation of this score (+1SD) shown by a dotted vertical line (x=0.31466). Variants with a score >ROC threshold but <+1SD are classed as having intermediate effect. Known oncogenic variants with intermediate effect are coloured grey, and GoF effect (>+1SD) shown in red. Variants below the ROC threshold are considered ‘normal’ (blue). Likely oncogenic variants coloured blue are presumed false negatives, gnomAD variants coloured blue are true negatives and act as a control. n=230 variants are stratified over ROC and +1SD thresholds, with variants showing a spectrum of k-means cluster 0 (i.e. enriched for NRF2-ontology mis-regulation) membership (as calculated in Fig.4e,i). **i)** Variants observed as part of the TRACERx NSCLC precision medicine programme p.W24C (HGVSc=ENST00000397062.8:c.72G>T, HGVSp=ENSP00000380252.3:p.Trp24Cys), and in a rare heterozygous germline syndrome p.T80K (HGVSc=ENST00000397062.8:c.239C>A, HGVSp=ENSP00000380252.3:p.Thr80Lys) are installed in genomes at nucleotide precision with Saturation-seq and are shown to be positively disruptive, with high specific disruption scores and Cluster 0 k-means membership.

We filtered TCGA for tumors with mutated *NFE2L2* within protein positions sampled in our Saturation-seq experiment (24:33 and 77:86). W24C, G31A and G81V variants were identified in ‘lung squamous cell carcinoma’ (LUSC) tumors with W24C and G81V also seen in a ‘head-neck squamous cell carcinoma’ (HNSC) and ‘kidney renal papillary cell carcinoma’ (KIRP) tumor, respectively; all samples had adjacent normal tissue (see Methods: ‘Correlation with somatic TCGA tumor data and germline variants’), allowing for the calculation of log-fold change in expression between tumor and normal tissue. L30del (inframe codon deletion, ‘uterine corpus endometrial carcinoma’ [UCEC]), E82G (KIRP), D29H (HNSC and ‘kidney renal clear cell carcinoma’ [KIRC]) tumors were also seen. LUSC and HNSC TCGA tumors are the canonical tissues for clinical NRF2 disruption and were confirmed to have significantly increased normalized mRNA counts for known NRF2 targets compared to their matched normal sample; with UCEC, KIRP and KIRC samples showing non-significant changes (**Extended Data Fig.4a**). In addition, many known NRF2 targets were found to be significantly changed by DESeq2 differential gene expression analysis of aggregated *NFE2L2* mutated LUSC tumor samples, compared to all normal TCGA LUSC tissue samples (**Extended Data Fig.4b**).

Importantly, statistically significant linear correlations (Pearson’s) were seen between Saturation-seq LFCs and TCGA *NFE2L2* mutant tumor matched normal LFCs for known NRF2 target genes (curated gene list [see Methods], filtered for those with a positive Saturation-seq log-fold change (LFC>0), **Fig.6a-d**). G31A (a recurrent, oncogenic, LUSC variant) has a significant, positive correlation (**Fig.6a**, R=0.68, p=8.3e-03), whereas a synonymous change at the same position G31G does not (**Fig.6b**, R=0.25, p=0.32), indicating that positive correlations are due to specific variant effects. Notably, W24C Saturation-seq LFCs are significantly, positively correlated with both W24C LUSC and HNSC tumor matched normal LFCs (**Fig.6c,d**). Weaker correlations were observed for G81V (KIRP) and L30del (UCEC) (R = 0.60, p = 0.0032 and R = 0.31, p = 0.054, respectively; **Extended Data Fig.4c**), while E82G (KIRP), D29H (KIRC, HNSC) and G81V (LUSC) were non-significant (p > 0.1 for all); disruption scores for G81V, L30del, E82G and D29H were 0.159, 0.918, 0.147 and −0.072, respectively. D29H appears to be a Saturation-seq false negative, likely due to low cell numbers for this variant (n=4, where the median across missense variants is n=10). G81V and E82G are likely true positives, supported by recurrent observation in LUSC (E82G in MSK-IMPACT) and positive disruption scores; lack of TCGA correlation may reflect limited gene detection and/or tissue context. Further to correlating changes in gene expression, we find that we can successfully report single-cell differential expression analyses for Saturation-seq derived transcriptomes, where we find that variants in both DLG and ETGE canonical motif regions show known and novel target mis-regulation (**Fig.6e,f**), with a range of computed specific disruption scores seen for missense variants in these regions (**Extended Data Fig.4d**). Saturation-seq differential expression data is presented in **Supplementary Table 2**.

### Saturation-seq is highly sensitive and specific, permitting somatic and germline *NFE2L2* variant scoring

Next, we sought to quantify the sensitivity, specificity and error rate of specific disruption scores in the distinction of known clinical, GoF missense/inframe deletion *NFE2L2* variants from germline variants in gnomAD (a healthy, population-ascertained cohort)^40^. We computed a ROC curve using our calculated specific disruption scores for 25 MSK-IMPACT ‘likely-/oncogenic’, GoF variants as true-positives (n=23 missense, n=2 inframe deletions), and 7 gnomAD variants as true negatives (expected to be non-compromising or likely-/benign, n=4 synonymous, n=2 missense, n=1 stop-gained). Variants were matched to scores with nucleotide-resolution using HGVSc identifiers. ROC analysis determines that the Saturation-seq specific disruption score for NRF2 is highly sensitive (84%) and specific (100%) at the ideal score threshold (0.02502), with a very high classification accuracy (>90%) (**Fig.6g**). The ideal specific disruption score – at which maximum sensitivity and specificity is achieved – is very close to zero (0.02502) as expected. ROC assessment is an important summary metric, however per variant interpretation in the assessment of classification validity is also important. To this end, we use the ideal specific disruption score (0.02502) and +1 standard deviation (SD) of this score to define variant effect classifications. On this basis, variants are classified as having either: a highly positive disruptive GoF variant effect (>1 SD of the ideal score), an intermediate variant effect (>ideal score and <1 SD of this score) or a normal variant effect (<ideal score).

Recurrent somatic variants identified in MSK-IMPACT span the full spectrum of disruption scores, with n=15/25 classified as GoF (highly positively disruptive) and n=6/25 as intermediate (moderately positively disruptive), n=4/25 are classed as having normal (non-disruptive) function (**Fig.6h**). Importantly, clustering of cells by transcriptome (which is pathway/target agnostic, **Fig.4e,i**) scales dynamically with specific disruption scores across classification thresholds, such that cells harbouring variants with intermediate effect scores are distributed across Clusters 0 and 1 (mean Cluster 0 membership = 13.9%), whereas cells harbouring variants with GoF level scores are seen predominantly in Cluster 0 (mean Cluster 0 membership = 86.6%) (**Fig.6h**). In contrast, gnomAD variants, which are germline alleles identified in a healthy population-cohort, have low specific disruption scores, clearly below the ROC threshold and are therefore classified as having normal NRF2 function; a mean of 1.9% of cells for these variants had Cluster 0 membership (**Fig.6h**).

*NFE2L2* is predominantly mutated in lung malignancies, we therefore searched data from the ‘TRAcking Cancer Evolution through therapy/Rx’ (TRACERx) non-small cell lung cancer (NSCLC) precision medicine programme where we identified a patient/tumor sample (CRUK0087) containing a Saturation-seq assessed variant W24C (HGVSc= ENST00000397062.8:c.72G>T, HGVSp=ENSP00000380252.3:p.Trp24Cys)^41^. W24C has a specific disruption score of 0.62, with 83% of cells in Cluster 0 (**Fig.6i**). As above, W24C is also seen in TCGA LUSC and HNSC samples, where NRF2 target misregulation in TCGA tissues is correlated with Saturation-seq detected misregulation (**Fig.6c,d**). Therefore, W24C demonstrates clear oncogenicity. Expanding on this, we searched TCGA LUSC and HNSC cohorts, identifying 109 accessions with *NFE2L2* mutations (98 samples had 1 mutation, 1 sample had 3 mutations and 4 samples had 2 mutations); this includes 97 missense variants (n=43 unique changes), together with 6 inframe deletions, 5 fusions and 1 frameshift insertion (all unique variants). Of note, n=29 missense mutations (n=68 cases) and 1 inframe codon deletion (n=1 case) are seen in oncogenic hotspot regions 24:33 and 77:86 scored by Saturation-seq. Of these we were able to compute disruption scores for 21 unique HGVSc variants in total, covering 18/29 missense mutations, with n=12/21 scored as GoF (and n=3/21 as intermediate and n=6/21 as normal).

*NFE2L2* mutations are largely found to be somatic, however rare cases of *de novo* heterozygous missense germline variants have been described^42^. These variants cause a multisystem disorder with immunodeficiency and neurological symptoms. White matter abnormalities, decreased homocysteine levels, cachexia, and increased glucose-6-phosphate dehydrogenase and increased glutathione reductase have been reported in these patients. Our dataset scored a reported missense *de novo* germline variant (T80K, HGVSc=ENST00000397062.8:c.239C>A, HGVSp=ENSP00000380252.3:p.Thr80Lys) which we found to be highly disruptive, indicating Saturation-seq can report the effects of germline and somatic variants for NRF2 (**Fig.6i**).

In summary, clinical summary metrics (**Fig.6g**) and per-variant analyses demonstrate that Saturation-seq is effective for clinical variant interpretation. The assay shows a false positive rate of 0 (1−specificity) and a false negative rate of 0.16 (1−sensitivity), indicating that false negatives are more common than false positives. This is expected given that the GoF classification integrates signals across multiple independent transcripts, making false positive calls unlikely. Accordingly, whereas clinically recurrent variants classified as normal should be interpreted with greater caution, GoF variants identified by Saturation-seq can be called with high confidence.

## Discussion

Saturation-seq resolves a fundamental limitation in variant functional genomics: the inability to assign rich, cell-resolved phenotypes to specific genomically installed variants at scale. By coupling CRISPR-based HDR editing in a barcoded haploid cell line with large scale droplet-based detection of genotype and whole transcriptome, the platform directly links endogenously installed variants to their transcriptional consequences in multiplex—without relying on fitness proxies, sgRNA sequences, or inferred genotypes.

Applied to *NFE2L2*, we demonstrate that our approach resolves the function of 230 variants across oncogenic hotspots in a single experiment. We compute disruption scores, derived from 82 curated NRF2 targets and see a bimodal distribution of functional change for missense variants, and a unimodal null distribution for synonymous variants. We find that scores map directly onto KEAP1 binding sites. Furthermore, distinct substitutions at the same residue can produce distinct transcriptional profiles—a granularity directly relevant to the allelic diversity uncovered by clinical sequencing^43^. Disruption scores separate known pathogenic from benign *NFE2L2* variants (in a curated truthset) with >90% accuracy where we observe a false positive rate of 0%, and a false negative rate of 16%. This asymmetry is informative: calling a variant as GoF requires coordinated mis-regulation across many independent transcripts, making erroneous positive calls inherently improbable. GoF calls can be made with high confidence while variants that score at or near thresholds deserve scrutiny rather than a straightforward benign classification. Notably, GoF truthsets remain limited by the relative scarcity of classified GoF variants in genomic databases compared to LoF; expanding these resources remains an important challenge for the field^44^. Beyond summary metrics, we score variants identified in a TRACERx NSCLC patient, TCGA tumor cohorts, and a rare germline *NFE2L2* syndrome case within the same experimental framework, Saturation-seq can therefore be useful for both oncology and rare disease studies and in both somatic and germline contexts^41,42^.

Of note, single-cell resolution, which is especially important for GoF loci, is a key strength of Saturation-seq. Operating cell-by-cell preserves quantitative variation in transcriptional mis-regulation and enables two complementary readout metrics: pathway-informed disruption scores and pathway-agnostic k-means clustering, which independently stratify variants (at least for NRF2), with Cluster 0 membership scaling predictably across the dynamic range of computed disruption scores. Variants with GoF-level scores concentrate in Cluster 0 (mean 86.6% membership across cells); those with intermediate scores distribute across clusters (mean 13.9%); importantly, gnomAD benign alleles remain almost entirely in Cluster 1 (mean 1.9%). Ontology analysis of Cluster 0 cells recovers NRF2 pathway components, glutathione metabolism, and reactive oxygen species responses without prior knowledge of these associations—an independent validation of biological specificity.

Several limitations are worth acknowledging directly. The haploid cell context used in this study precludes assessment of allele-dosage effects relevant to heterozygous variants; however, our genotyping approach is amenable to diploid/triploid and heterozygous cells^24^. HAP1 cells faithfully recapitulate canonical NRF2 responses however exceptions may exist in non-canonical tissues. Per-variant differential expression is constrained by cell numbers (median n=10 per missense variant); the D29H false negative, for which barcodes and variants were detected in n=4 cells illustrates that additional depth would improve sensitivity. Cell dropout during bottlenecking reduced variant recovery to roughly 57% of the post-bottleneck pool; this is directly addressable by retaining more cells through the bottlenecking step. In terms of NRF2 targeting, disruption scores are calibrated to a curated 82-gene target set, so neomorphic effects on non-canonical programmes are not captured by the score alone, though pathway-agnostic clustering and ontology enrichment partially mitigate this.

Our approach changes how variant effect mapping can be approached for GoF loci at scale, indeed as variants are installed within their genomic context, non-coding/intronic/promoter/UTR/splicing variants could also be investigated. SGE with fitness readouts has proven transformative for essential tumor suppressors, where pathogenic/disruptive variants manifest as viability phenotypes readily quantifiable by amplicon sequencing across thousands of variants. Oncogenes present a fundamentally different problem: pathogenic variants often do not compromise fitness but instead rewire transcriptional programmes whose complexity, and genotype-phenotype correlation, is lost in bulk approaches. Saturation-seq addresses this by directly linking genotype to phenotype and reallocating experimental resources—from broad fitness-based coverage of thousands of variants to deep single-cell phenotyping of hundreds within discrete, saturated target regions—and by replacing fitness with transcriptomic readouts. The framework generalises to any gene for which a tractable transcriptional phenotype can be defined, with particular promise for transcription factors, signalling regulators, and oncogenes, classes most underserved by existing interpretation tools. Ideal targets combine a validated cell context with well-characterised downstream targets to maximise the dynamic range for disruption scoring. However, as demonstrated here, pathway-agnostic clustering of single-cell transcriptomes can stratify variants by molecular phenotype even where downstream programmes are incompletely characterised, broadening the approach to less well-understood loci. Furthermore, stop-gained and frameshift variants yield negative disruption scores, consistent with NRF2 LoF—notable given GoF is the predominant mutational/pathological mechanism—suggesting that Saturation-seq can capture transcriptomic responses at non-essential LoF loci, broadening applicability across genetic contexts. Additionally, as these data comprise unbiased single-cell whole transcriptomes for >200 variants, they constitute a resource for further query without the need for additional experiments, e.g. nucleotide-resolution process/structure comparisons.

Key developments could expand and improve the Saturation-seq framework. The adoption of prime editing—despite current limitations in editing efficiency—would allow many genomic loci to be saturated simultaneously, while emerging joint DNA-RNA profiling platforms would streamline single-cell genotyping and phenotyping, increase cell throughput, and limit cellular stress—improving cell recovery, library complexity, and assay sensitivity^45–48^. Together, these advances would deepen single-cell differential expression resolution, enabling variant-specific transcriptional responses to be mapped under defined conditions such as hypoxia or drug treatment, and grading target gene responses in ways directly relevant to therapeutic investigation. Complementing transcriptomics with single-cell epigenomic and proteomic readouts would extend the technology further still, capturing variant effects across molecular layers currently beyond reach—broadening Saturation-seq into a multimodal platform for variant interpretation in personalised medicine^49–52^.

## Methods

### Clonal line production – HDR template construction

HAP1-A5 cells were edited to introduce specific genomic variants. HDR templates were generated using PCR Site Directed Mutagenesis (SDM) of a wild-type genomic region *E.coli* cloned into the pMin plasmid (NEB Q5® Site-Directed Mutagenesis Kit, following manufacturer’s instructions)^4^. In detail, genomic DNA (gDNA) was isolated from the HAP1-A5 cell line using the Qiagen DNeasy Blood & Tissue Kit, following the manufacturer’s protocol. PCR amplification was carried out with KAPA HiFi HotStart ReadyMix–using primers that amplify a wild-type genomic region that included the DLG and ETGE regions in addition to 5’ appending sequence complementary to vector backbone sequence (primers ‘clonal_line_HDR_F/R’ were used, **Supplementary Table 3**)–and 100ng of HAP1-A5 genomic DNA as template in a 50 µL reaction. Reactions were run for fewer than 30 cycles, using an annealing temperature of approximately 65 °C and a 90 sec extension step at 72 °C. PCR products were analyzed on 0.8% agarose-TAE gels to confirm single, correctly sized bands, which were then purified using the Qiagen QIAquick Gel Extraction Kit. A plasmid vector fragment containing the cloning adapters, origin of replication, and ampicillin resistance cassette was similarly amplified from the ‘pMin-U6-ccdb-hPGK-puro’ backbone using ‘hr_frag_f/r’ primers (see **Supplementary Table 3**). The PCR was performed with KAPA HiFi HotStart ReadyMix at 63 °C annealing for fewer than 25 cycles, followed by DpnI digestion to remove template plasmid. The expected 1.8 kb product was separated on a 0.8% agarose-TAE gel and purified using the QIAquick Gel Extraction Kit. Assembly of wild-type genomic amplicons with the vector fragment was carried out using NEBuilder® HiFi DNA Assembly Master Mix (NEB). Fragments were combined at a 2:1 molar ratio (50 ng vector fragment) and incubated for 1 h at 50 °C in a thermocycler. The assembled reaction was diluted 1:4 with water, and 2 µL of the diluted mixture was transformed into 50 µL of TOP10 competent *E. coli* (Invitrogen). Transformants were plated on LB agar containing 100 µg/mL ampicillin and incubated overnight at 37 °C. Individual colonies were picked for liquid culture and plasmid isolation using the QIAprep Spin Miniprep Kit. Insert sequences were verified by Sanger sequencing (Eurofins) using ‘hdr_seq_f/r’ primers (**Supplementary Table 3**).

A confirmed wild-type region clone was used as template for subsequent site-directed mutagenesis generation, which used the NEB Q5® Site-Directed Mutagenesis (SDM) kit (following manufacturer’s protocol) and using SDM prefix primers listed in **Supplementary Table 3**. Bacterial clones were selected, grown in Ampicillin (100 µg mL⁻¹) LB culture, mini-prepped (Qiagen) and the desired plasmid sequence confirmed by Sanger Sequencing using primers ‘neh2_hdr_min_f/r’ (**Supplementary Table 3**). Glycerol stocks were expanded and maxi-prepped (Qiagen). sgRNAs for clonal line generation were cloned as described in Methods: ‘sgRNA cloning’ using the same sequences as for the main Saturation-seq experiment.

### Clonal line production – genome editing

HAP1-A5 cells were thawed and expanded as in Methods: ‘Saturation Genome Editing of NRF2 target regions’. 1e+6 cells were seeded into 12 well plates (Corning) in 1mL Iscove’s Modified Dulbecco’s Medium (IMDM, Gibco) 10% Foetal Bovine Serum (FBS, Gibco) 1% Penicillin/Streptomycin media (Gibco) one day before transfection (day-1). On the day of transfection (day 0), culture medium was replaced one hour before transfection. Transient transfection was performed using Xfect (Takara) according to the manufacturer’s instructions. Xfect buffer was thawed from −20 °C at 4 °C overnight and equilibrated to room temperature before use. For each transfection, 1 µg of sgRNA plasmid and 2 µg of SDM generated HDR template plasmid were combined in an Eppendorf tube, and Xfect buffer was added to a total volume of 50 µL. Freshly thawed Xfect polymer (1.8 µL; 0.6 µL per µg DNA) was then added, followed by vortexing and a 10 min incubation at room temperature. Transfection mixes were vortexed and 50 µL of the mixture was added dropwise to each well of a 12 well plate. Cells were incubated with this mix for 4 h at 37°C, after which the medium was replaced with fresh medium and cultures were incubated overnight.

On days 1 and 2 post-transfection, the medium was replaced with selection medium containing blasticidin (10 µg mL⁻¹) and puromycin (3 µg mL⁻¹; InvivoGen) to maintain *Cas9* expression and select for transfected sgRNA plasmids. On day 3, cells were split 1:2 into new 12 well plates in blasticidin-containing medium and incubated overnight. On day 9, cells were split 1:2 into 6 well plates in IMDM 10%FBS 1% P/S media (no selective antibiotics). Polyclonal cell populations were banked in liquid nitrogen on day 18; this included the following lines DLGΔ, E79Q, T80K, G31R and D29* (KEAP1Δ obtained via Horizon Bioscience, was also single cell sorted).

Polyclonal vials were thawed and expanded over 7-10 days in T75 flasks in IMDM 10%FBS 1% P/S media, with 5e+6 cells seeded at each passage. 2e+6 cells were suspended in media in FACS test tubes, with Hoechst added to final concentrations of 10 µg mL⁻¹ (for ploidy gating, selecting haploid cells). Cells were incubated for 15mins at RT. Cells were centrifuged at 300rpm for 10mins, washed with 5mL of PBS and resuspended in FACS buffer containing Propidium Iodide (PI, to select for alive cells) at a final concentration of 1 µg mL⁻¹. Cells were sorted into single wells of 96 well plates in 180 µL of IMDM 10%FBS 1% P/S media. Non-stained controls were included for gating. Cells were cultured and expanded at day 14 (post-FACS) were passaged into 24 well plates, with 12 clones selected per variant line. Plates were cloned. One plate was kept in culture, another frozen at −80°C. DNA extractions were performed on cells in culture, and PCRs on each extraction using ‘neh2_gdamp_f/r’ (**Supplementary Table 3**) using 350ng of gDNA in 50 µL PCR reactions, which were subsequently purified using a PCR clean-up kit (Qiagen, 40 µL EB elution), run on 0.8% agarose gel to confirm successful PCR, and Sanger sequenced (Eurofins) in 96 well plates, using primers ‘neh2_seq_f/r’ (**Supplementary Table 3**). Sanger chromatograms with single mutant peaks at intended edited sites were recorded as confirmed clones, and frozen samples for these clones, thawed, cultured and expanded into T75 flasks and re-banked in bulk.

### sgRNA cloning

sgRNA target sequence oligonucleotides were synthesized with overhangs for directional cloning: 5ʹ-CACC-3ʹ was added to the sense strand (or 5ʹ-CACCG-3ʹ if the target sequence did not begin with G, to optimize U6 promoter transcription) and 5ʹ-AAAC-3ʹ to the antisense strand (with an additional 3ʹ-C when using CACCG on the sense strand). Each oligonucleotide (1 µL at 100 µM) was phosphorylated and annealed in a 10 µL reaction containing 0.5 µL polynucleotide kinase (PNK), 1 µL 10× T4 ligation buffer (NEB), and nuclease-free water. Reactions were incubated at 37 °C for 30 min, then gradually cooled from 95 °C to 25 °C at a rate of 5 °C per minute. Annealed oligonucleotides were diluted 1:200 prior to ligation.

For vector preparation, 20 µg of maxi-prepped ‘pMin-U6-ccdb-hPGK-puro’ (pMin) plasmid was digested with BbsI (NEB) in a 100 µL reaction using 10 µL 10× CutSmart Buffer (NEB) for 3 h at 37 °C. The 3,653 bp backbone was gel-purified using a 0.8% agarose–TAE gel, divided between two wells, and cleaned using the QIAquick Gel Extraction Kit (Qiagen), followed by a MinElute purification (Qiagen). DNA purity and concentration were verified via NanoDrop.

Ligation was performed by combining 1 µL of diluted annealed oligonucleotides with 50 ng of gel-purified pMin backbone, 5 µL 2× Quick Ligase Buffer (NEB), and 1 µL Quick Ligase (NEB), incubating at room temperature for 10 min. The ligation mix was diluted 1:4 with water, and 2 µL was transformed into 50 µL of *E. coli* TOP10 competent cells (Invitrogen). Colonies were selected on ampicillin-containing (100 µg mL⁻¹) LB agar, inoculated into LB broth with ampicillin, and cultured overnight. Glycerol stocks were prepared, and correct sgRNA inserts were confirmed by Sanger sequencing (Eurofins) using ‘guide_seq_f/r’ primers (**Supplementary Table 3**). Verified sgRNA plasmids were expanded in 125 mL LB broth with ampicillin and purified by maxiprep (Qiagen) to obtain transfection-grade DNA.

### Western blots

Cells were washed with ice-cold PBS, then scraped in 1mL PBS and transferred to 1.5mL Eppendorf tubes and centrifuged (13,000rpm 5 minutes at 4°C) and supernatant removed. Cells were then lysed in ice-cold RIPA lysis buffer (CST) containing EDTA-free protease inhibitor cocktail (Merck) with regular vortexing over 30 minutes with incubation on ice. The solution was centrifuged (13,000rpm 5 minutes at 4°C) and protein-containing supernatant transferred to a new 1.5mL Eppendorf tube. Protein concentration was determined using a BCA assay kit (ThermoFisher) in technical triplicate. Equal amounts (5µg) of protein per sample were separated by SDS–PAGE gel in MOPS buffer (Invitrogen) with 0.25 % antioxidant solution (180V for 45 minutes). Gels were then transferred to PVDF membranes in transfer buffer (Invitrogen). Membranes were stained with Ponceau and blocked with 5% milk in TBST for 1 hour at RT. Blocked membranes were incubated with primary antibodies against NRF2 (Abcam, Rabbit monoclonal, AB62352, 1:1,000 dilution), NQO1 (Promega, goat polyclonal, AB2346, 1:10,000 dilution), KEAP1 (Sigma, MABS514/Clone 144, 1:1000 dilution) and GAPDH (Cell Signalling Technology, 14C10/2118S, rabbit monoclonal, 1:1,000 dilution) overnight at 4°C. After washing 3x with TBST, membranes were incubated with secondary antibodies–either Anti-goat, donkey polyclonal (Abcam, AB97110) at 1:2,000 or Anti-rabbit, goat polyclonal (Abcam, AB97051) at 1:2,000–for 1 hour at RT and then sprayed with WesternBright ECL (Advansta, K-12049-D50), and signals captured on a developer.

### Colony forming assay

Cells were seeded at a density of 800 cells per dish in 5mL Iscove’s Modified Dulbecco’s Medium (IMDM, Gibco) 10% Foetal Bovine Serum (FBS, Gibco) 1% Penicillin/Streptomycin media (Gibco) in 100mm petri dishes and cultured for 7 days under standard conditions (37°C, 5% CO₂). After incubation, dishes were washed with PBS, colonies were fixed and stained with 5mL of 20% methanol and 0.5% crystal violet solution (Sigma) for 30 minutes at room temperature (RT). Excess stain was washed off with tap water, and plates were air-dried overnight. Dried Petri dishes were photographed (with black and white filter) and colony number compared.

### MTS and Almar Blue assays

Cell viability was assessed using MTS (Promega, G5421) and Alamar Blue (ThermoFisher, A50100) assays according to the manufacturer’s instructions. Cells were seeded in 96 well plates at a density of 5,000 cells per well (in 100μl IMDM) and treated with Cisplatin (Selleckchem, S116614) over a concentration range of 0 to 8µM. For the MTS assay, 40 µL of prepared MTS reagent (as per manufacturer’s instructions) was added to each well and absorbance was measured at 490nm and 630nm each hour for 4 hours in total using a plate reader. For the Alamar Blue assay, 20 µL of prepared reagent (as per manufacturer’s instructions) was added to each well and fluorescence was measured at 570nm and 600nm wavelengths each hour for 4 hours in total using a plate reader. Results were normalized to untreated wells. Results were analysed in GraphPad Prism where, by default, a logistic dose-response model is fit by nonlinear least squares, with asymmetric 95% confidence intervals derived by profile likelihood.

### Clonal line RT-qPCR

RNA was extracted from clonal cell lines (both genome edited lines and the KEAP1Δ commercial line) and HAP1-A5 control using a RNAeasy mini kit (Qiagen) together with QIAshdredder homogenization (β-mercaptoethanol was added to RLT buffer), from 0.5e+6 cells seeded into 6 well plates and cultured for four days, RNA was eluted in 30μL RNAase free water. RNA concentration and purity were assessed by Nanodrop spectrophotometry, and diluted to 50ng/μl. One step RT-qPCR (i.e. gene specific reverse transcription followed by cDNA quantitation in one reaction) was performed using a Luna® Universal One-Step RT-qPCR Kit, with FAM-MGB probes for genes of interest (*AKR1C1*, *NQO1*, *GCLM*, *GCLC* and *NRF2*) and VIC-MGB probe for the endogenous housekeeping gene 18s. RT-qPCR was run on low ROX machine (Applied Biosystems, QuantStudio 7 Flex). Reactions were performed in technical triplicate, with non-template controls per gene. Relative gene expression was calculated using the ΔΔCt method, normalized to 18s, and expressed relative to HAP1-A5.

DLGΔ and E79Q clonal cell lines were assayed for continued elevated canonical target gene expression (*NQO1* and *AKR1C1*) over cell culturing/time to ensure molecular phenotypes persist beyond immediate editing events. The same RNA extraction and RT-qPCR process was followed for these experiments, except that three biological replicates rather than technical replicates were used (except for *AKR1C1* timepoints D4 and D10 in both lines, where one biological replicate was run).

### Clonal line single cell sequencing

HAP1 cells were dissociated using TrpLE (ThermoFisher), and the cell suspension was passed through a 30μm cell strainer. Cells were washed in PBS with 0.04% BSA and loaded onto the 10x Genomics 5’ Next-Gem Chip K with the aim to recover 4,000 cells per lane (1 lane per clonal line). Sequencing libraries were generated following the manufacturer’s protocol for 5’ NextGem v2 kits, and cells were sequenced with a depth of at least 50,000 reads per cell.

### Cloning of transcribed cell barcode library in a lentivector

The U6 promoter and guide scaffold were first removed from lentivector (Addgene #67974), by cutting with MluI and BamH1, treating with Klenow (NEB) to produce blunt ends, and relighting the vector. To introduce the cell barcode (in the 5’ UTR of the BFP transcript), a single stranded ultramer containing NeoUTR3 was amplified using KAPA to add Gibson arms and a 10N barcode in the reverse primer (see **Supplementary Table 3** for sequences)^53^. After SPRI purification, the product was cloned by Gibson (NEB) into the new lentivector (without the U6 promoter/guide scaffold) cut with XhoI and BsrGI (which also removed the puro selectable marker). After ethanol precipitation, 8 Gibson reactions were electroporated into 8 aliquots of super competent *E.coli* cells (Endura, Lucigen), pooled and grown in liquid culture to give a coverage of around 200 million barcodes (quantified by serial dilutions of the culture spread onto LB plates and colony counted). A maxi prep (Qiagen) of the cell barcode plasmid library was made from the cell pellets. A 250 bp PCR product covering the cell barcode was generated (see **Supplementary Table 3** for sequences) using the plasmid library as a template, and amplicon sequencing was performed using Miseq to confirm that there was no overrepresentation of any of the barcode sequences (see Methods: ‘Next generation sequencing of edited genomes and HDR template libraries’).

### Lentivirus production and HAP1 cell barcoding

The cell barcode library was transfected into 239FT cells in Opti-MEM (Gibco), using lipofectamine LTX (Invitrogen), along with the packaging/envelope plasmids psPax2 (Addgene #12260) and pMD2.G (Addgene #12259) and the virus was harvested 48hrs later. Virus was transduced into dissociated HAP1 cells using 8 µg/mL polybrene and titrated to give an MOI of less than 0.3 using flow cytometry to measure BFP expression. A large-scale transduction of HAP1-A5 cells constitutively expressing Cas9 and containing a 10bp deletion in *LIGASE IV* (*LIG4*) and sorted for 1N haploidy (previously optimized for high SGE editing rates and Cas9 activity, Horizon HZGHC-LIG4-Cas9), was performed to maintain coverage of the cell barcode library^4^. To sort both BFP positive cells and haploid cells, a control cell line known to be haploid was stained with Hoechst (10μg/mL) for 1hr at 37°C before performing flow cytometry to gate cells with 1N DNA content (haploid cells, as HAP1 cells can revert to diploidy in culture at low levels)^54^. These 1N gates were transferred to an FSC/SSC plot to allow sorting of haploid cells that were non-Hoescht stained and also BFP positive. A total of 8 million cells 1N BFP cells were sorted (in 4 batches) from an original transduction of 4-5 million cells (18 million cells with an MOI of 0.3). See **Extended Data Fig.1** for representative FACS plots and gating conditions.

### SGE library design

HDR template library sequences were generated *in silico* using VaLiAnT (and subsequently, unique oligonucleotide sequences were chemically synthesized by Twist Bioscience, see Methods: ‘HDR template library construction’)^37^. All sequences were based on the GRCh38 genome build, with coordinates chosen to define the DLG and ETGE regions limits for variant introduction as follows: Chr2:177234162-177234281 and Chr2:177233983-177234122, for DLG and ETGE regions, respectively. These ranges were defined by liminal primer sequences: F_TTCTGTTTTTCCAGCTCATACT, R_TTAAACATAGGACATGGATTTGA and F_ CCAAGAACTGAGTACTCTGTACCT, R_ GCAAGAGAAAGCCTTTTTCGC, for DLG and ETGE regions, respectively. Systematic VaLiAnT mutator functions were used to generate a selection of mutation categories in ranges within the target region defined by the liminal primers; these sub-ranges were: Chr2:177234218-177234247 and Chr2:177234059-177234088, which correspond to the peptide ranges W24-R25-Q26-D27-I28-D29-L30-G31-V32-S33 and D77-E78-E79-T80-G81-E82-F83-L84-P85-I86, for DLG and ETGE target regions, respectively.

The mutator functions were inputted as a vector into VaLiAnT (version 2.0.0, https://github.com/cancerit/VaLiAnT) and were as follows: ‘snvre’ which generates all SNVs and all possible synonymous changes at codons, ‘aa’ which generates all missense changes at codons using the default VaLiAnT codon usage table (most common codon sequences for *homo sapiens*), ‘1del’ which installs frameshifting 1bp deletions, ‘stop’ which installs stop-gained mutations and ‘inframe’ which introduces in frame triplet codon deletions across the defined peptide ranges (see above). Custom variants with sequential deletions of codons within DLG and ETGE were also generated. The *in silico* generation produced n=423 (n=333 unique) DLG region variants and n=425 (n=329 unique) ETGE region variants.

sgRNAs (TGGAGGCAAGATATAGATCT, WGE ID:942562361 and GTTACAACTAGATGAAGAGAC, WGE ID: 942562354, for DLG and ETGE, respectively) were selected within each defined range based on criteria outlined in Barbon *et al*.^37,55^. Fixed synonymous changes were included at the PAM/Protospacer regions for all oligonucleotides generated to prevent Cas9 re-cutting installed DNA tracts; we term these synonymous changes ‘PAM/Protospacer Protection Edits’ (PPEs). Therefore, all oligonucleotide sequences generated contain a variant and a PPE. PPEs were as follows: 177234236_A>G, 177234239_T>C and 177234242_C>T for sgRNA-942562361 and 177234095_T>C, 177234092_T>C, and 177234089_T>C for sgRNA-942562354.

Mission Bio primers for single-cell amplicon sequencing over the DLG and ETGE combined region within the Neh2 domain of NRF2 (‘Mission_Bio_NRF2_F/R’, **Supplementary Table 3**) were designed with a proprietary algorithm to work with an array of other primers to produce a multiplexed amplicon library (including over the 10N barcode) which has sufficient DNA to permit sequencing.

Accurate chemical synthesis of oligonucleotides is limited to roughly <300nt, therefore generated oligo pools, must be appended 5’ and 3’ with genomic sequence extensions that permit HDR, theses extensions are termed homology arms. Homology arm tracts were defined using primers either side of the DLG and ETGE target regions and such that one Mission Bio amplicon sequencing primer was outside this range to obviate single-cell amplification of HDR template library (see **Supplementary Table 3**).

### HDR template library construction

The starting wild-type sequences (NRF2 target regions with suitable arms of homology, sequences in **Supplementary Table 3**) for each library were synthesised by Twist Biosciences and cloned into the basic pTwist Amp High Copy vector: https://www.twistbioscience.com/products/genes/vectors?tab=catalog-vectors).

For each target region, the homology arm and variant library oligo lengths used were as follows. For the DLG target region, the 5’ homology arm was 165bp, the variant library oligo 120bp, and the 3’ homology arm 800bp. For the ETGE target region, the 5’ homology arm was 800bp, the variant library oligo 140bp, and the 3’ homology arm 218bp. The wild-type plasmids were linearised by PCR (adding Gibson arms) (see ‘PlasmidLin’ sequences, **Supplementary Table 3**), while at the same time removing the wild-type target region sequence to be replaced by with variant library. The variant libraries were ordered from Twist Biosciences and amplified by PCR to yield a pool of double-stranded DNA with ‘Gibson assembly’ cloning arms (see **Supplementary Table 3** for oligo sequences)^56^. The variant libraries were cloned into the linearised homology arm plasmids by Gibson assembly (NEB) with at least 4 Gibson reactions for each target region to maintain coverage. After ethanol precipitation, each Gibson reaction was electroporated into an aliquot of super competent *E.coli* cells (Endura, Lucigen), pooled per target region and grown in liquid culture to give at least 1000x coverage of each library (quantified by serial dilutions of the culture spread onto LB plates and colony counting). A maxi prep (Qiagen) of each library was made from the cell pellets, and the plasmid libraries were sequenced using amplicon sequencing on a Miseq to check for accurate coverage of all variants (Methods: ‘Next generation sequencing of edited genomes and HDR template libraries’).

sgRNA sequences were cloned into the minimal expression vector pMin as above (Methods: ‘sgRNA cloning’).

### Saturation Genome Editing of NRF2 target regions

Vials containing approximately 5e+6 HAP1-A5 barcoded cells were thawed and seeded into T75 tissue culture flasks (Corning) nine days prior to transfection in 15 ml of medium supplemented with blasticidin (10 µg mL⁻¹; InvivoGen) to select for cells carrying an integrated *Cas9* construct. Cells were passaged at a 1:10 ratio six days before transfection and expanded three days prior into multiple T150 flasks containing 35 mL of blasticidin-supplemented medium.

One day before transfection, a cell stock suspension was prepared by resuspending 8e+6 cells in 15 mL of medium without blasticidin. A 15 mL aliquot of this suspension was seeded into each T75 flask designated for transfection. On the day of transfection (day 0), culture medium was replaced one hour before transfection.

Transient transfection was performed in duplicate (replicates were used for next generation sequencing QC, Saturation-seq pools were derived from single transfection populations) using Xfect (Takara) according to the manufacturer’s instructions. Xfect buffer was thawed from −20 °C at 4 °C overnight and equilibrated to room temperature before use. For each transfection, 7.5 µg of sgRNA plasmid and 15 µg of HDR library plasmid were combined in an Eppendorf tube, and Xfect buffer was added to a total volume of 750 µL. Freshly thawed Xfect polymer (13.5 µL; 0.6 µL per µg DNA) was then added, followed by vortexing and a 10 min incubation at room temperature. Replicate transfection mixes were pooled, vortexed, and 750 µL of the mixture was added dropwise to each T75 flask. Cells were incubated for 4 h at 37 °C, after which the medium was replaced with fresh medium and cultures were incubated overnight.

On days 1 and 2 post-transfection, the medium was replaced with selection medium containing blasticidin (10 µg mL⁻¹) and puromycin (3 µg mL⁻¹; InvivoGen) to maintain *Cas9* expression and select for transfected sgRNA plasmids. On day 3, cells were split 1:2 into a T75 flask in 15 mL of blasticidin-containing medium and incubated overnight. On day 4, each T75 flask was bottlenecked and passaged and the remainder of cells harvested for genomic DNA collection. Cells were washed once with 5 mL PBS (Sigma), dissociated using 1.5 mL TrypLE Express (no phenol red; ThermoFisher) at 37 °C for 4 min, and neutralized with 1.5 mL medium lacking antibiotics. The cell suspension was transferred to a 15 mL Falcon tube, and the flask was rinsed with an additional 5 mL of medium. Viable cell counts were determined using a Countess Automated Cell Counter (ThermoFisher) with trypan blue staining (Gibco). To reduce barcode complexity and see the same barcode-variant combination represented in single-cell DNA and single-cell RNA sequencing, the edited cell populations were bottlenecked at 3,000 cells per transfected T75 flask (we required ∼10 barcodes per variant, with a total of ∼300 variants per target region = 3000 cells) and seeded into wells of 96 well plates in 150 µL of media. The remaining cells were pelleted at 300 g for 3 min, washed once with 1 mL PBS, and centrifuged again. Pellets were resuspended in PBS at a final concentration of 6 × 10⁶ cells mL⁻¹, aliquoted (1 mL per tube), centrifuged at 300 g for 3 min, and stored at −80 °C until genomic DNA extraction (this formed pre-bottlenecked samples for next generation sequencing). On day 8, 100% of bottlenecked cell populations were seeded into 12 well plates and were expanded into T75 flasks on day 11. On Day 14, multiple vials of 6e+6 cells per transfection were pelleted and frozen in IMDM, 20% FBS, 10%DMSO media, and transferred to −80 °C and subsequently into liquid nitrogen storage. Vials were also pelleted (as above) for a post-bottlenecking next generation sequencing samples.

### Next generation sequencing of edited genomes and HDR template libraries

Genomic DNA (gDNA) was extracted from pre-bottlenecked and post-bottlenecked pelleted sample cell pellets using the Qiagen DNeasy Blood and Tissue Kit, following the spin-column protocol for cultured cells. Prior to lysis, samples were treated with RNase A (Qiagen) for 15 minutes at 37°C to remove RNA contamination. DNA was eluted in 100 µL of AE buffer, and concentration and purity were assessed using a Nanodrop spectrophotometer. Only samples with concentrations greater than 50 ng/µL, A260/A280 ratios between 1.8–1.9, and A260/A230 ratios above 2 were used for subsequent PCR analyses. Samples that did not meet these quality criteria were re-extracted.

Sequencing libraries were prepared through three PCR steps using KAPA HiFi HotStart ReadyMix (Roche) in 50 µL reactions with 0.3 µM primers: a primary PCR to enrich edited loci, a secondary PCR to add Illumina adapters, and an indexing PCR for sample labelling. For the primary PCR, 4 µg gDNA (∼1e+6 haploid genomes) was split into four 1 µg reactions to maintain edited gDNA library complexity. One primer was placed outside the homology arm to avoid amplifying residual HDR plasmid (primers ‘gdamp_f/r’; **Supplementary Table 3**), producing 1.2–1.9 kb amplicons. Reaction conditions were pre-optimized identifying annealing at 63°C for 23 cycles. Four reactions were pooled and purified with QIAquick columns (Qiagen), adding 3 µL NaOAc to PB buffer to adjust pH, and eluted in 40 µL EB buffer. To remove primers, products were treated with Exonuclease I (NEB) (5 µL enzyme, 10 µL buffer, 37°C 20 min; 80°C 20 min), purified with MinElute (Qiagen), and eluted in 15 µL EB. Concentration and purity were checked by Nanodrop, and products were diluted to 10 ng/µL for the secondary PCR.

Secondary PCRs, using ‘DLG_HDR_Library_Amplicon F/R’ or ‘ETGE_HDR_Library _Amplicon F/R’ primers (**Supplementary Table 3**), appended Illumina adapters, generating DLG (119 bp), and ETGE (137 bp) amplicons. Optimal conditions were 63°C annealing and 15 cycles. 75 ng of primary product was used as template across three reactions, which were pooled and purified as above.

HDR plasmid libraries (DNA HDR template containing variants used in SGE editing) were amplified using 100 ng plasmid DNA divided into four 50 µL reactions, with ‘DLG_HDR_Library_Amplicon F/R’ or ‘ETGE_HDR_Library _Amplicon F/R’ primers (**Supplementary Table 3**) and the same cycling conditions as the primary PCR. Products were purified as above and diluted to 5 ng/µL for indexing.

Indexing PCRs used 25 ng secondary PCR template, annealed at 59°C for 8 cycles with Illumina D501–D504 (i5) and iPCRtag (i7) primers. Products were purified using AMPure XP beads (0.9× ratio), eluted in 33 µL EB, and normalized to 10 ng/µL by Nanodrop. 15 µL of each library was prepared; 8 µL was run on a 2% agarose–TAE gel (120 V, 45 min) to confirm a single band, and 5 µL was pooled per target region. Pooled libraries (10 ng/µL) were quantified by KAPA qPCR and Agilent TapeStation, diluted to 8 pM, and sequenced with 20% PhiX spike-in. gDNA libraries and plasmid libraries were sequenced on an Illumina MiSeq v3 (150-cycle, PE) platform run and analysed using the QUANTS pipeline (See GitHub Repository, and below).

### Informatics Processing and Editing Quantification

Raw sequencing data in CRAM format were processed using the QUANTS pipeline (version 1.2.1.0; https://github.com/cancerit/QUANTS/releases/tag/1.2.1.0) to generate sequencing quality control metrics and perform exact-match mapping of reads to the designed VaLiAnT library sequences. QUANTS (https://github.com/cancerit/QUANTS) is a Nextflow-based pipeline built using the nf-core framework^57^. Analyses were conducted with Nextflow version 22.04.3 and Singularity, which provided the containerized software environment. QUANTS version 1.2.10 was executed using three primary inputs: (i) raw CRAM sequencing files, (ii) the HDR template library reference, and (iii) a sample mapping file linking each sequencing file to a defined sample identifier.

Within QUANTS, adapter trimming was performed using cutadapt (version 3.2, Python 3.8.6) with the parameters --cores 4 -a [adapterR1]…[adapterR2]^58^. Quality control (QC) metrics and plots were generated for both raw and trimmed reads using FastQC (version 0.11.9; https://github.com/s-andrews/fastqc) and SeqKit (version 0.15.0; stats function), with all outputs aggregated via MultiQC (version 1.10.1)^59,60^. Library-independent read quantification was carried out using pyCROQUET (version 1.5.0; https://github.com/cancerit/pycroquet), which enumerated the frequency of each unique trimmed read sequence. These counts were then processed using a custom R script included within the QUANTS repository (https://github.com/cancerit/QUANTS/tree/1.2.1.0/modules/local/R/post_pycroquet_quantification) to determine the abundance of each HDR template. Reads not mapping to designed library sequences were also quantified. To estimate editing efficiency, reads mapping to the ref_seq (wild-type GRCh38 reference) and pam_seq (wild-type sequence containing only PAM/protospacer protection edits) were compared.

### Single cell sequencing for genotyping (Mission Bio) of transcriptomics (10x Genomics)

Both pools (DLG and ETGE) of bottlenecked edited cells were thawed and expanded to T75 flasks and single cells were processed as for the clonal lines above. (see Methods: ‘Clonal line single cell sequencing’). The cells were single-cell genotyped using the Tapestri machine (Mission Bio, v3 kit, according to manufacturer’s instructions), one reaction for each edited pool of cells (DLG and ETGE), with up to 15,000 cells per reaction and using a custom panel of amplicon sequences (including primers to amplify the BFP cell barcode, and primers to amplify around the NRF2 exon – see **Supplementary Table 3** for sequences). The same population of cells was also loaded onto the 10x Genomics 5’ HT Chip, one lane for each edited pool of cells (DLG and ETGE), with the aim to recover 60,000 cells per lane. We spiked a BFP transcript specific RT primer (2 µL of 10 µM stock, ‘BFP_RT Ultamer’ see **Supplementary Table 3** for sequence) into the cell lysis mixture (made up according to manufacturer’s protocol) before GEM formation, in order to capture the transcribed BFP cell barcode specify and create an enriched library)^24^. After the cDNA amplification stage, we performed a 0.6x SPRI cleanup (to recover the large DNA fragments containing the full-length cDNA) and then a further 1.3X SPRI cleanup to recover the small DNA fragments (containing the shorter cell barcode transcript). We performed nested PCR from both the small and the large SPRI fractions to produce separate libraries for sequencing the BFP cell barcodes (see **Supplementary Table 3** for primer sequences). cDNA libraries were prepared according to the manufacturer’s instructions from the large SPRI fraction. Sequencing was performed on 250PE Miseq (Illumina) for the Mission Bio genotyping libraries and on the NovaSeq 6000 (Illumina) for the 10x Genomics and cell barcode libraries.

### Analysis of clonal scRNA-seq data sets

Cell Ranger v. 7.0.1 was used to obtain UMI counts as well as for cell calling. Cells with low total UMI count, low outliers for the number of detected features and high outliers for the percentage of UMI counts from mitochondrial genes were identified using the scater Bioconductor package and discarded, retaining 25,823 cells^61^. Differential expression was performed using the negative binomial approach implemented in the NEBULA Bioconductor package^62^.

### Saturation-seq DNA bioinformatics analysis

Reads were mapped to a reference of amplicons using BWA-MEM with a quality score cutoff of 30^38^. Droplets with at least 2 filtered reads each for the *NFE2L2* and UTR-barcode amplicons were retained. We then discarded reads not matching either the wild-type genotype or a successful SGE outcome, e.g. reads with indels or editing outcomes other than those defined by SGE. To remove empty droplets, doublets, diploid cells and noise reads, any genotype-droplet combinations that did not constitute at least 90% of the reads for the droplet were discarded, resulting in 1,927 and 1,928 droplets for the DLG and ETGE data sets respectively. For each droplet, if there was not one UTR-barcode that covered at least 90% of the reads, the Hamming distance between the UTR-barcodes that together make up at least 90% of the reads for the droplet was computed. For distances >1, the droplet was discarded. For distances = 1, the UTR-barcode with the highest number of reads was assigned.

UTR-barcode-genotype combinations with less than 3 cells were discarded. This resulted in 217 and 242 unique barcodes and 149 and 154 unique genotypes for the DLG and ETGE data set respectively, including wild-type cells (with PPE variants, to normalize for editing effects, as above) and those with successful SGE outcomes.

### Saturation-seq RNA processing and UTR-barcode calling

Cell Ranger v. 7.0.1 was used to obtain UMI counts for both mRNA and UTR-barcodes, as well as for cell calling. Cells low for the total UMI count, low outliers for the number of detected features and high outliers for the percentage of UMI counts from mitochondrial genes were identified using the scater Bioconductor package and discarded, leading to 55,872/56,783 cells for the DLG/ETGE data sets respectively^61^.

### Saturation-seq barcode, variant and transcriptome linkage

A list of UTR-barcodes present in the DNA data was compiled from the Saturation-seq DNA bioinformatics analysis. From the RNA data, these UTR-barcodes were called using a method applying mixtures of skewed normal distributions to robustly assign barcodes and gRNAs to cells^24^. This identified 11,379/11,440 cells for the DLG/ETGE data sets for which we were able to confidently call an UTR-barcode present in the DNA data. 1,853/1,917 out of these cells had one of the 217/242 UTR-barcodes confidently associated with the SGE or wild-type (PPE containing) outcomes for the DLG/ETGE data sets (see Methods: ‘Saturation-seq DNA bioinformatics analysis’).

### Differential expression analysis using mixed effects models

Differential expression analysis between the wild-type and the SGE-edited genotypes was performed using generalised linear mixed models (GLMMs) as implemented in the nebula R package^62^. This allowed capturing the hierarchical structure of genotypes being represented by multiple different groups of cells sharing the same UTR-barcode in terms of a mixed effects model. Multiple testing adjustments were performed using the standard Benjamini-Hochberg method^63^.

### Integrated and specific disruption score computation

‘Integrated disruption scores’ were obtained by first integrating the DLG and ETGE data sets based on 82 known targets of *NFE2L2* derived from an expert review, using the mutual nearest neighbours method, and then computing the first principal component (PC) of the integrated dataset^29,64^. ‘Specific disruption scores’ are equal to the first principal components obtained from each of the two data sets individually, again based on the known *NFE2L2* targets. Differences in integrated disruption scores across variant consequences were tested for using the Mann-Whitney U-test.To annotate variant metadata, VaLiAnT design outputs (version 2.0.0) in VCF format were used to compute VEP (via online portal, March 2024) annotations, and merged with score dataframes based on VaLiAnT variant ID^37,65^. The variant Chr2:177234244_T>A was filtered from score dataframes; this variant falls into a codon with a PPE variant (AGG>AGA), leading to a dinucleotide change (AGA>TGA) upon systematic SNV generation which results in a stop-gained mutation rather than a missense change of R25W, as annotated by VEP.

### Cell-based clustering of transcriptomes and ontology analysis

We applied K-means clustering (spanning 2 to 5 clusters, random seed = 42) to the log-transformed counts for DLG (1853 cells, 385 genes) and ETGE (1917 cells, 249 genes). Based on silhouette score evaluation, the 2-cluster model performed best for both DLG (0.0517) and ETGE (0.0799) and was selected for downstream analysis (**Fig.4e,i**). Next, we performed differential expression analysis between the two clusters utilizing DESeq2 on raw count data^66^. Significantly differentially expressed genes (adjusted p < 0.01) were extracted and analyzed for pathway enrichment using the ‘Express Analysis’ in Metascape^67^.

### Correlation with somatic TCGA tumor data and germline variants

For matched-normal tumor tissue comparisons, pan-cancer TCGA batch-normalized mRNA expression data were downloaded from the UCSC Xena Browser on January 29, 2025 (dataset ID = ‘EB++AdjustPANCAN_IlluminaHiSeq_RNASeqV2.geneExp.xena’, version 2016-12-29). The corresponding TCGA clinical metadata were obtained from the same source and date (dataset ID = ‘Survival_SupplementalTable_S1_20171025_xena_sp’, version 2018-09-13). Samples harbouring reported variants in *NFE2L2* (NRF2) were identified by searching the pan-cancer TCGA dataset within the Xena Browser. During data integration, a small number of samples (TCGA-19-5956-01, TCGA-BG-A220-01, TCGA-BT-A42B-01, TCGA-EY-A1G8-01, and TCGA-FZ-5924-01) were found to be absent from the mRNA expression dataset and were excluded from downstream analyses. A known issue in the TCGA mRNA dataset involves a duplicate entry for *SLC35E2*, as previously reported (https://github.com/hamidghaedi/Methylation_Analysis); this duplication was resolved prior to analysis. Only samples derived from solid tissue (TCGA code “11” denoting solid tissue normal) were retained, while blood-derived normal samples were excluded due to their limited relevance for RNA-based analyses.

Filtering of the curated mutation list found six variants (with matched normal samples) identified in *NFE2L2* that were screened by Saturation-seq: G31A, G81V, W24C, L30del, E82G and D29H. These corresponded to the following TCGA tumor samples/tissues contexts: G31A_LUSC_tumor = TCGA.22.5483.01, G81V_LUSC_tumor = TCGA.51.4079.01, W24C_LUSC_tumor = TCGA.56.7582.01, L30del_UCEC_tumor = TCGA.BG.A3EW.01, E82G_KIRP_tumor = TCGA.BQ.7044.01, G81V_KIRP_tumor = TCGA.BQ.7051.01, W24C_HNSC_tumor = TCGA.CV.6935.01, D29H_HNSC_tumor = TCGA.CV.7424.01, and D29H_KIRC_tumor = TCGA.CZ.5456.01. Solid tissue matched normal samples were as follows: G31A_LUSC_normal = TCGA.22.5483.11, G81V_LUSC_normal = TCGA.51.4079.11, W24C_LUSC_normal = TCGA.56.7582.11, L30del_UCEC_normal = TCGA.BG.A3EW.11, E82G_KIRP_normal = TCGA.BQ.7044.11, G81V_KIRP_normal = TCGA.BQ.7051.11, W24C_HNSC_normal = TCGA.CV.6935.11, D29H_HNSC_normal = TCGA.CV.7424.11, and D29H_KIRC_normal = TCGA.CZ.5456.11.

For DESeq2 analyses (shown in **Extended Data Fig.4b**) TCGA LUSC samples harbouring single missense somatic mutations within the DLG or ETGE motifs of *NFE2L2* that were recurrently seen (>2 tumors) were identified and filtered using the UCSC Xena Browser (February 2025)^66,68^. All TCGA LUSC normal solid tissue samples were also filtered. *SLC35E2* duplicate accessions were removed as above. Raw RNA-Seq read count data (RSEM expected counts) were downloaded from the UCSC Xena Browser (dataset ID = ‘tcga_gene_expected_count’, version 2016-09-01, downloaded February 2025) to ensure unnormalized count input for DESeq2 (counts were rounded to integers using 2^x^, where x=RSEM count). DESeq2 analysis was performed on TCGA tumors which contained *NFE2L2* variants, each tumor sample was considered a biological replicate (total n=31; D29H=4, D29N=5, D29Y=2, E79Q=7, G31A=5, G81S=2, L30F=4, T80K=2) for DESeq2 experimental condition design. Solid tissue LUSC matched normal samples for filtered recurrent variants were used as control samples (n=3); (i.e. specific pairwise analyses were not performed due to lack of tumor/normal sample extraction replicates). DESeq2 (version DESeq2_1.46.0) was run and FDR (BH adjusted p-values) used to define up (FDR<0.01, LFC>0), down (FDR<0.01, LFC<0) and non-significant change (FDR>0.01) in gene expression. The following known targets were highlighted in **Extended Data Fig.4b** if significantly mis-regulated (these are also the targets used to define disruption scores); *AKR1B10*, *AKR1C1*, *AKR1C2*, *AKR1C3*, *ALDH1A1*, *ALDH7A1*, *CBR1*, *CES1H*, *EPHX1*, *NQO1*, *PTGR1*, *GSTA1*, *GSTA3*, *GSTM1*, *GSTP1*, *MGST1*, *SULT1A1*, *UGT1A6*, *UGT2B7*, *ABCB6*, *ABCC1*, *ABCC2*, *ABCC3*, *CAT*, *GPX2*, *GPX4*, *PRDX1*, *PRDX6*, *BLVRB*, *FECH*, *FTH1*, *FTL*, *HMOX1*, *SLC40A1*, *GCLC*, *GCLM*, *GGT1*, *SLC7A11*, *TXNRD1*, *SRXN1*, *G6PD*, *PGD*, *IDH1*, *ME1*, *TALDO1*, *TKT*, *CD36*, *MARCO*, *PSMA1*, *PSMA4*, *PSMA7*, *PSMB1*, *PSMB2*, *PSMB3*, *PSMB5*, *PSMB6*, *PSMC1*, *PSMC3*, *PSMD1*, *PSMD5*, *PSMD11*, *PSMD12*, *PSMD13*, *PSME1*, *SQSTM1*, *CALCOCO2*, *ULK1*, *ATG7*, *GABARAPL1*, *LAMP2*, *AHR*, *ATF3*, *ATF4*, *CEBPB*, *MAFG*, *PPARG*, *PPARGC1B*, *RXRA*, *KEAP1*, *NFE2L2*, *TRPA1* and *NOTCH1*^29^.

## Data and Code

All bioinformatic and analysis code has been deposited at: https://github.com/StatOVarI-lab/Saturation-seq

## Acknowledgements

This work was supported by the Cancer Research UK CG-MAVE programme (EDDPGM-Nov22/100004). The Wellcome Sanger Institute is supported by core funding from the Wellcome Trust (grant 220540/Z/20/A). Additional support for barcode design and implementation was provided by Open Targets (OTAR2071). The authors would like to thank Dr. Sebastian Gerety, Dr. Katrina Andrews, and members of the Atlas of Variant Effects Alliance (https://www.varianteffect.org/) for useful discussions.

---

**Extended Figure 1:**
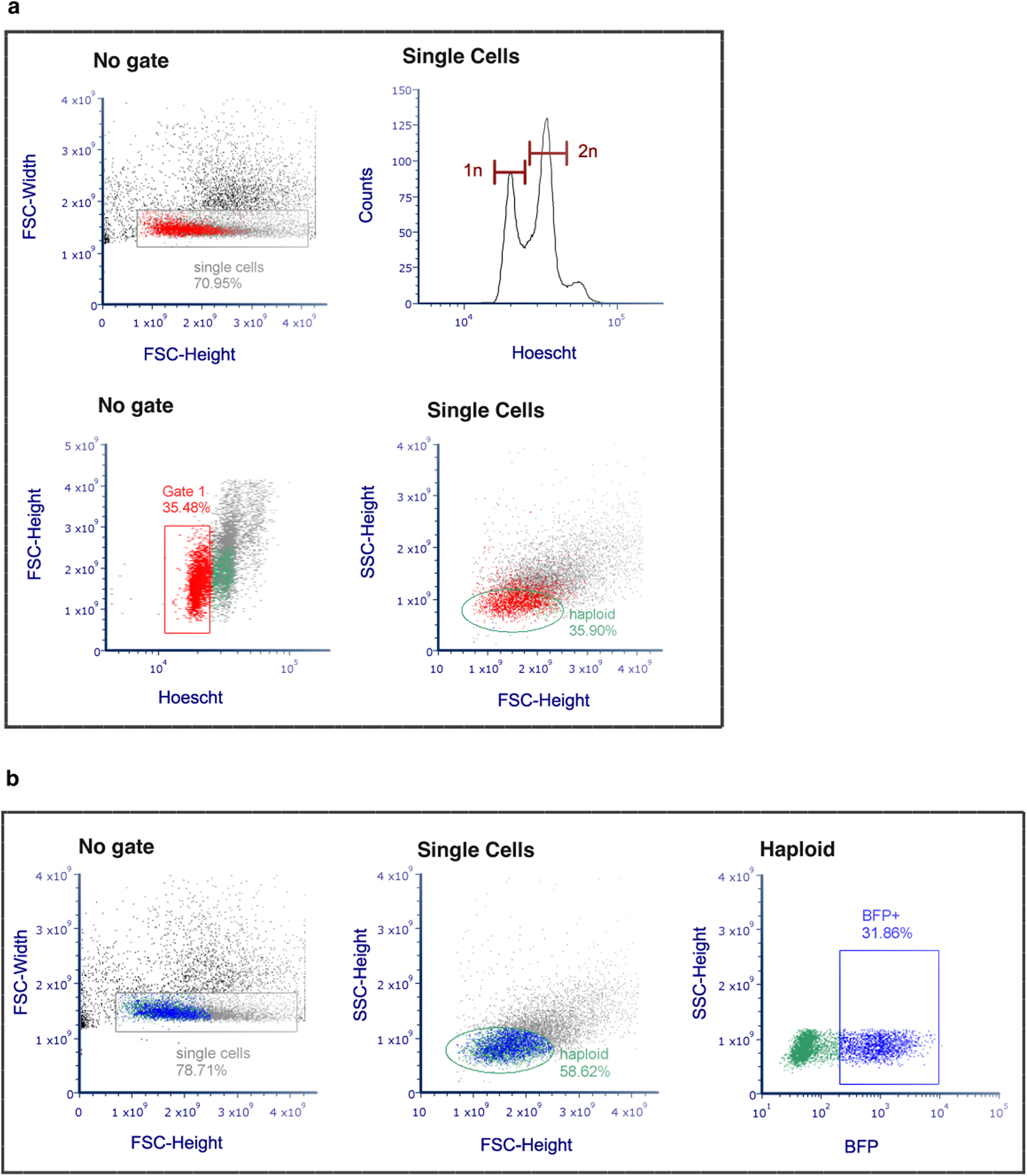
FACS gating for haploidy and BFP+ barcode transduction sorting. **a)** Haploid control cells can be sorted based on size. True haploid cells were identified using Hoescht staining for 1n DNA content and the gates were transferred to an FSC/SSC plot to allow sorting of haploid cells that were non-Hoescht stained. **b)** Haploid barcode expressing cells were sorted. Using the same FSC/SSC gates, haploid cells that were also BFP positive were sorted. These cells could not be Hoescht stained for haploid 1n sorting, as the BFP fluorophore would be masked. Data showing batch 1 of the sorting. Identical gates were used for 4 batches which resulted in a total of 8 million cells sorted to generate a complex pool of cell barcodes.

**Extended Figure 2:**
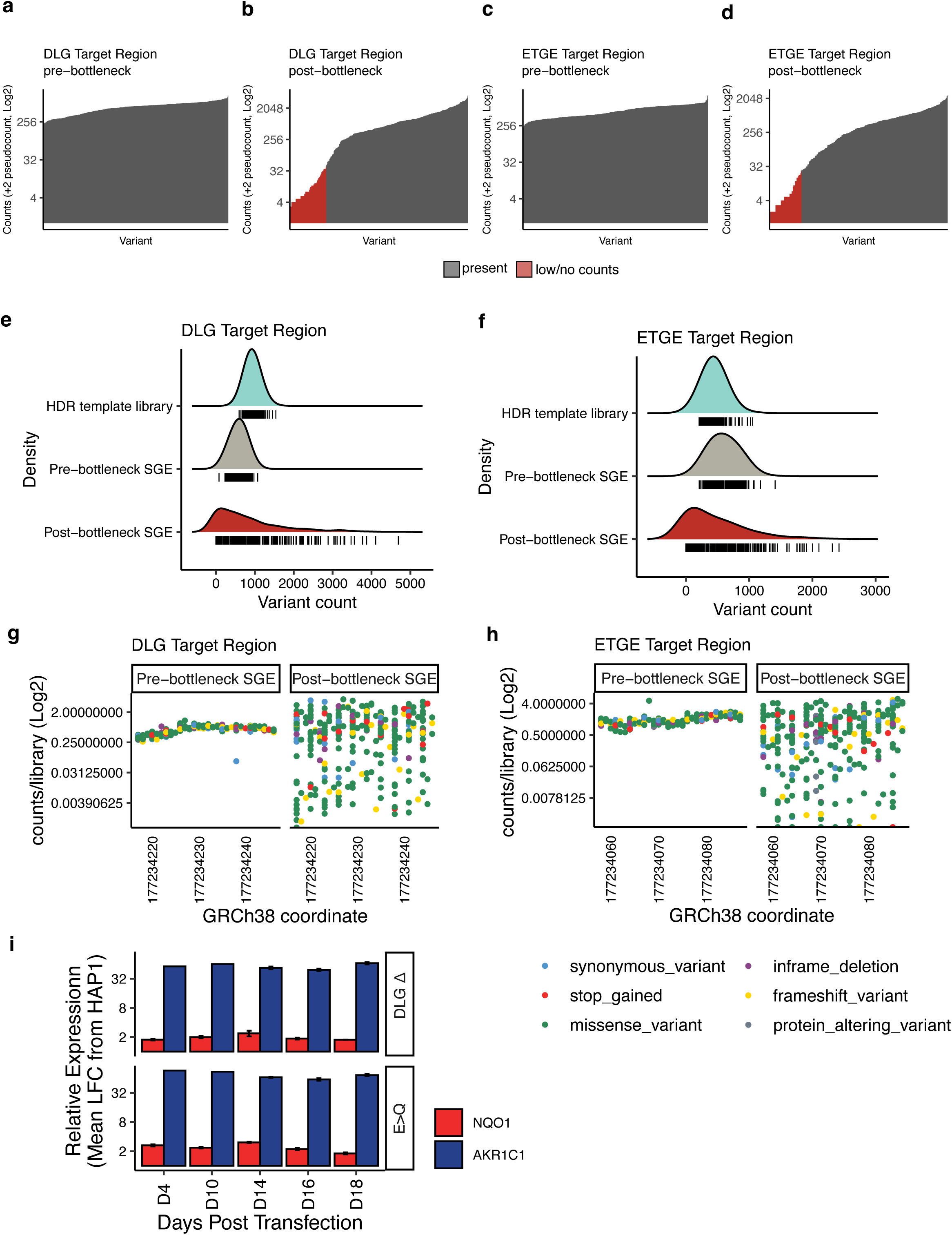
Editing efficiency, bottlenecking and time-point validation. **a)** Mapped counts (y axis) per variant (x axis) from a pool of pre-bottlenecked cells for DLG variants, all variants are well represented in the edited pool. **b)** As ‘a’ but reads for post-bottlenecked DLG cells (3,000 cells seeded and expanded), low/no counts variants (<5% of expected proportion of mapped counts, i.e. 100%/333 or 329 variants = ∼0.3%*0.05 cutoff, therefore variants at <0.015% representation) are highlighted in red. **c-d)** as ‘a’ and ‘b’ but for ETGE region, respectively. **e)** Variant count density distributions for HDR template library plasmid (light blue), and saturation edited genomes before bottlenecking (grey) show normal distributions of variant representation, after bottlenecking (red), the distribution changes to a positively skewed binomial distribution, expected of sampling for biological count data under a selective pressure (bottlenecking); this confirms variant count representation with limited loss of variant representation. **f)** as for ‘e’ but with ETGE data. **g-h)** Genome editing positional effects are limited, for DLG (g) and ETGE (h) loci, as measured by pre-bottleneck counts as a ratio of HDR template library plasmid counts, coloured by VEP mutational consequence. Post-bottlenecked cells are not biased by consequence representation. **i)** Clonal line (DLGΔ and E79Q) qPCR data demonstrating that target gene expression (*NQO1* and *AKR1C1*) remains high from early timepoints, until after Day 14, the terminal screening timepoint employed (see **Fig.3b**) (geometric mean of 3 biological replicates, except D4 and D10 *AKR1C1*, 1 sample), validating that GoF molecular misfunctions likely persist until scRNA-Seq stages, allowing phenotype capture.

**Extended Figure 3:**
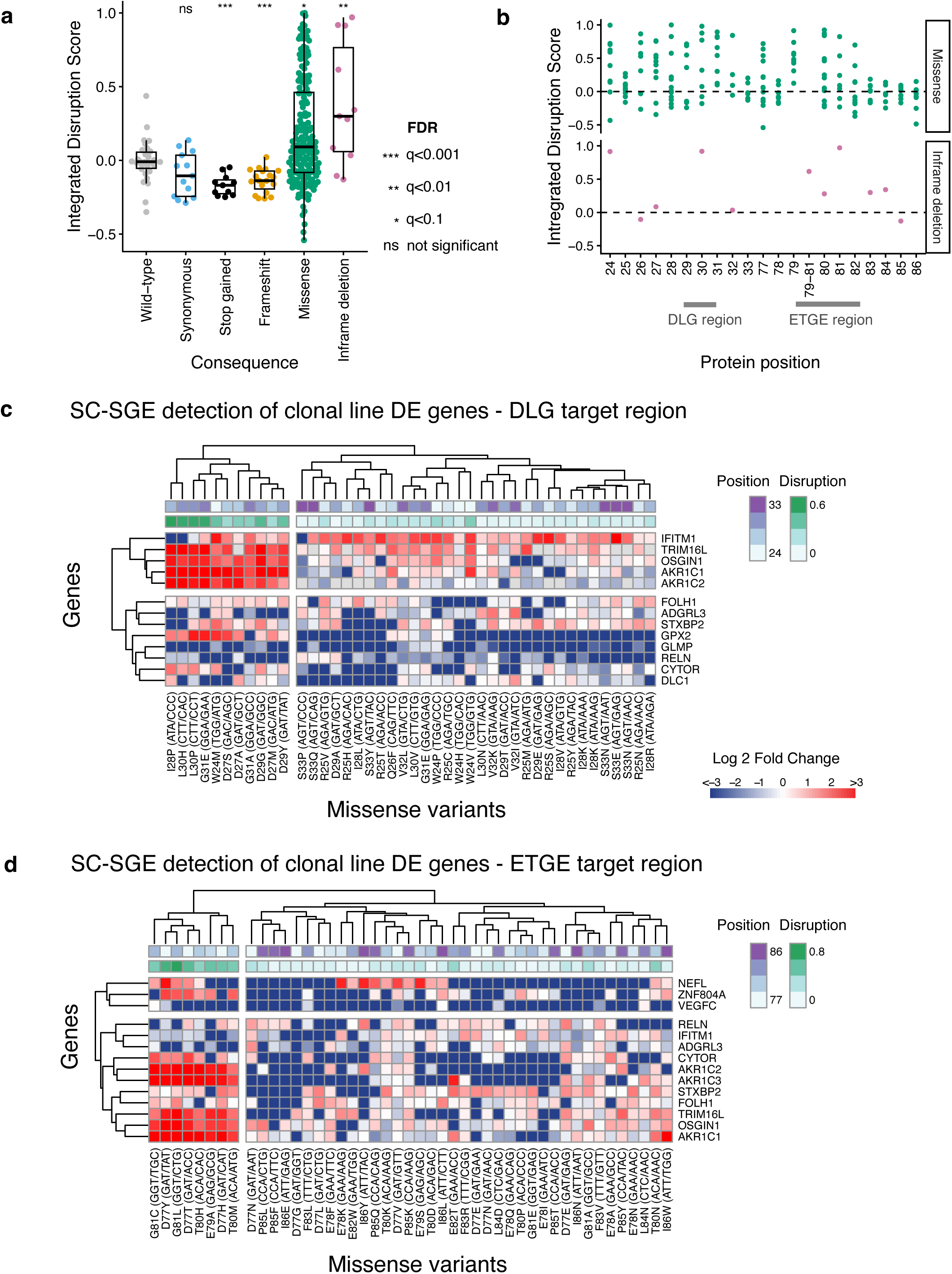
Mutational consequence disruption and clonal line differential gene expression at single-cell level. **a)** Variants stratified across mutational consequences show significantly different median disruption scores compared to edited wild-type; missense variants (green) show a broad spectrum of change with a significantly positive difference to wild-type. Stop-gained and frameshift show significantly negative disruption in aggregate, inframe codon deletions show two clearly distinct clusters which are in aggregate significant. Synonymous changes are not significantly different from wild-type. Mann-Whitney U test BH FDR q value thresholds are shown. **b)** The distribution of variant effect across protein positions shows that specific missense variants show diversity of disruption, with 83-86 showing little disruption across multiple variants (top panel, green). Inframe deletions show distinctly increased disruption within the cores of the DLG and ETGE motifs and at the recurrently mutated W24 codon (bottom panel, pink). **c)** Heatmap coloured by LFC from wild-type (red for increase, blue for decrease) for those genes previously found to be differentially expressed (DE) in clonal line studies in the DLG target region. Key NRF2 target genes cluster into a distinct over-expression cluster (top left) and are associated with variants in certain regions of the protein (purple) and with higher disruption scores as expected (green). Codon triplet changes leading to missense changes are shown across the x-axis. **d)** As ‘e’ but for ETGE target region and with the over-expression cluster at the bottom left.

**Extended Figure 4:**
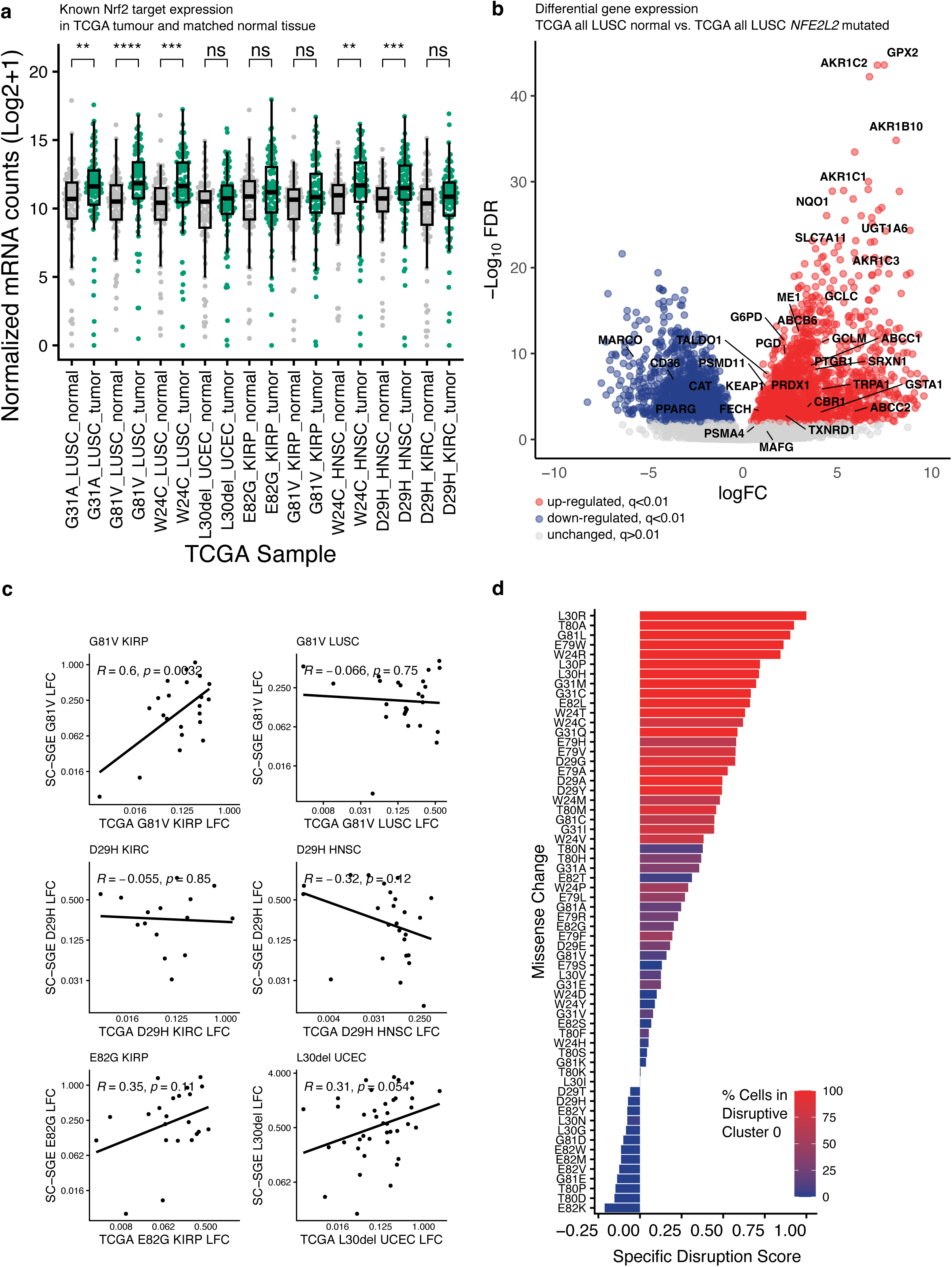
TCGA tumor RNA-seq analysis confirms known NRF2 targets are mis-regulated in lung and head and neck cancers. **a)** Nine *NFE2L2* mutant tumor samples with matched normal tissue (non-blood) are available in TCGA, namely G31A, G81V, W24C (LUSC), L30del (UCEC), E82G, G81V (KIRP), W24C, D29H (HNSC), D29H (KIRC). RNA counts for 82 known targets of NRF2 are plotted for tumor samples (green) and normal samples (grey). In aggregate, known targets (median) are significantly increased in expression compared to cognate matched normal samples in tumors/tissues known to be recurrently mutated at *NFE2L2* (LUSC and HNSC), but not in other tissues. Mann-Whitney U test, **** p<0.0001, *** p<0.001, ** p<0.01, ns=not significant. Boxes show the interquartile range, the horizontal lines show the median *mRNA* count and whiskers show the maximum and minimum values that are not outliers. **b)** All TCGA LUSC normal samples (n=119) were grouped and mRNA expression compared to all TCGA LUSC samples (grouped, n=31) in which *NFE2L2* was called as mutated (single missense) within DLG or ETGE regions, regardless of matched-normal status. A volcano plot representing differential expression analysis via DESeq2 on these two groups of samples is shown (red points represent up-regulated genes, blue down-regulated genes and grey unchanged expression). Known targets that are significantly up or down regulated are labelled. **c)** SC-SGE LFC comparisons to TCGA tumor LFCs for matched-normal samples/variants. G81V and E82G are moderately disruptive (scores 0.159 and 0.147) but not recurrently mutated in their respective TCGA tissues (KIRP/LUSC; KIRP). L30del is highly disruptive (score 0.918) but not recurrent in UCEC. D29H is scored as non-disruptive (score −0.072) and recurrent in HNSC but not in the KIRC sample shown. **d)** Missense variants screened by saturation-seq at the canonical KEAP1 binding positions in DLG 29:31 and ETGE 79:82 show a spectrum of specific disruption scores, cells of variants with positive scores are predominately found in Cluster 0 by ‘*de novo’* (i.e. known target agnostic) dimensionality reduction of transcriptomic effect (See Fig.4e,h,i,l), conversely variants that are scored as non-disruptive/LoF are predominately found in Cluster 1, as expected.

